# Evaluation of epigenetic age acceleration scores and their associations with CVD related phenotypes in a population cohort

**DOI:** 10.1101/2022.07.06.498980

**Authors:** Olga Chervova, Elizabeth Chernysheva, Kseniia Panteleeva, Tyas Arum Widayati, Natalie Hrbkova, Jadesada Schneider, Vladimir Maximov, Andrew Ryabikov, Taavi Tillmann, Hynek Pikhart, Martin Bobak, Vitaly Voloshin, Sofia Malyutina, Stephan Beck

## Abstract

We evaluated associations between nine epigenetic age acceleration (EAA) scores and 18 cardio-metabolic phenotypes using an Eastern European ageing population cohort richly annotated for a diverse set of phenotypes (subsample, n = 306; aged 45-69 years). This was implemented by splitting the data into groups with positive and negative EAAs. We observed strong association between all epigenetic age acceleration scores and sex, suggesting that any analysis of EAAs should be adjusted by sex. We found that some sex-adjusted EAA scores were significantly associated with several phenotypes such as blood levels of gamma-glutamyl transferase and low-density lipoprotein, smoking status, annual alcohol consumption, multiple carotid plaques, and incident coronary heart disease status (not necessarily the same phenotypes for different EAAs). We demonstrated that even after adjusting EAAs for sex, EAA-phenotype associations remain sex-specific, which should be taken into account in any downstream analysis involving EAAs. The obtained results suggest that in some EAA-phenotype associations, negative EAA scores (i.e. epigenetic age below chronological age) indicated more harmful phenotype values, which is counter-intuitive. Among all considered epigenetic clocks, GrimAge was significantly associated with more phenotypes than any other EA scores in this Russian sample.

## 1. Introduction

It has been more than a decade since the very first epigenetic age predictor was proposed [1], and since then dozens of DNA methylation (DNAm) based clocks have been developed. “Epigenetic age” (EA) is a score that is calculated by applying an EA prediction model (a DNAm clock) onto a set of DNA methylation measurements at particular loci (CpGs). Epigenetic age acceleration (EAA) is defined as the deviation of the estimated EA from the chronological age (CA), and is typically derived as either the difference between EA and CA, or as the residual from regressing EA onto CA (EA ~ CA).

In the beginning of the epigenetic clock era, the first generation EA predictors (i.e. [1], [2], [3]) were primarily focused on accurate age prediction. The new EAA measures, which are derived from second generation epigenetic clocks (i.e. [4], [5]), are more focused on capturing physiological dysregulation [6] while still keeping strong links to chronological age [7].

Various measures of EAA are shown to be associated with different phenotypes and diseases (see reviews [8] and [9]). For example, deviations in EAA were shown to be connected to cancer [10,11], metabolic syndrome [12], and cognitive function decline [13]. All of these conditions are linked with ageing, which is a complex process that involves changes in all organs, tissues, and cells; and cannot be quantified by a single biological measure. Similarly, there is no single EAA measure that could be declared as the best epigenetic marker of ageing.

In this study we investigate the relationship between several widely-used EAA scores with the phenotypic data on cardio-vascular disease (CVD) related risk factors and conditions available for a random population sample (*n* = 306) that is a part of The HAPIEE Project [14], a Siberian cohort established in 2003 as a multicentre epidemiological study of CVD in Eastern and Central Europe. One of our aims was to determine which EAA measures are “sensitive” to which phenotypes and health-related conditions. By comparing the distributions of phenotypes between those with positive and negative EAAs we identified how EAAs are associated with clinical data in an ageing Russian population.

## 2. Methods

### 2.1. Data collection

This study is based on the data generated from a subset of the Russian branch of the HAPIEE (Health, Alcohol, and Psychosocial Factors in Eastern Europe) cohort [14], which was established in Novosibirsk (Russia) in 2003-2005 and followed up in 2006-2008 and then again in 2015-2017. The protocol of the baseline cohort examination included an assessment of cardiovascular and other chronic disease history, lifestyle habits and general health, socio-economic circumstances, an objective measurement of blood pressure (BP), anthropometric parameters, physical performance, and instrumental measurement. The details of protocol are reported elsewhere [14].

This study is based on a cohort of *n* = 306 HAPIEE participants who did not have any indications of cardiovascular disease during the baseline measurement, as well as having a whole blood DNA methylation profile. DNAm was measured in accordance with the manufacturer’s recommended procedures using the Illumina MethylationEPIC BeadChip (Illumina, San Diego, CA, USA), a detailed description is available in [15]

### 2.2. Variables description

All variables involved in our analyses were collected during the baseline examination (with the exception of incident coronary heart disease). Phenotypic data available for our study includes age, sex, systolic and diastolic blood pressure values (SBP and DBP, mmHg; respectively), anthropometric parameters - body mass index (BMI, kg/m^2^) and waist-hip ratio (WHR, units), smoking status (ever smoker or never smoker), and the estimated annual alcohol intake (g of ethanol and number of annual occasions). A person who smoked at least one cigarette a day was classified as an “ever smoker”. The amount of alcohol consumed was assessed using the Graduated Frequency Questionnaire and was then converted to pure ethanol (g) [16]. The height and weight was measured with accuracy to 1 mm and 100 g, respectively. Blood pressure (BP) was measured three times (Omron M-5 tonometer) on the right arm in a sitting position after a 5 minute rest period with 2 minutes interval between measurements. The average of three BP measurements was calculated and recorded.

Fasting blood serum tests’ results contain measured levels of total cholesterol (TC, mmol/l), triglycerides (TG, mmol/l), high-density lipoprotein cholesterol (HDL, mmol/l), gamma-glutamyl transferase (GGT, mmol/l) and plasma glucose (mmol/l). The levels of TC, TG, HDL, GGT, and glucose in blood serum were measured enzymatically with the KoneLab 300i autoanalyser (Thermo Fisher Scientific Inc., USA) using Thermo Fisher Scientific kits. The Friedewald formula [17] was applied to calculate low-density lipoprotein cholesterol (LDL, mmol/l). Fasting plasma glucose (FPG) was calculated from the fasting serum glucose levels using the European Association for the Study of Diabetes (EASD) formula [18]. Hypertension (HT) comprises SBP ≥ 140 mmHg or DBP ≥ 90 mmHg according to the European Society of Cardiology/European Society of Hypertension (ESC/ESH) Guidelines [19] and/or antihypertensive medication intake within two weeks prior to the blood draw. Presence of type 2 diabetes mellitus (T2DM) was defined as FPG > 7.0 mmol/l, or ongoing treatment with insulin or oral hypoglycaemic medicines [20]. None of the participants included in our analysis had a history of major cardiovascular disease (CVD), such as myocardial infarction (MI), acute coronary syndrome (ACS), stroke, or transit ischemic attack at the time of the baseline examination and blood draw. The binary coronary heart disease (CHD) variable in our dataset includes any incident CHD events (MI/ACS) which occurred within the 15-year follow-up period of the cohort.

Carotid arteries were examined via high resolution ultrasound using the systems Vivid q or Vivid7 (GE HealthCare) with a 7.5/10-mHz phased-array linear transducer. Device settings were adjusted in accordance to the American Society of Echocardiography (ASE) recommendations [21]. Longitudinal and transverse scans were performed at the right and left common carotid arteries with branches to assess anatomy and atherosclerotic lesions. The digital images were archived and the measurements were conducted off-line by an experienced researcher (A.R.) who was blinded to the participants’ characteristics [22]. The plaques were defined in accordance with the Mannheim consensus [23]. For the present analysis we used two phenotypes of atherosclerosis: presence of at least one carotid plaque (CP) or multiple plaques (MCP). The ultrasound variables are only available for a subset of samples (*n* = 105, 35% of all samples).

Individual phenotypes were also combined into five groups of phenotypes, which we define as follows:

1. **Anthropometric**: BMI and WHR;
2. **Lifestyle**: smoking status and annual alcohol consumption (intake and number of occasions);
3. **Metabolic**: GGT, T2DM and plasma glucose;
4. **Lipids**: TC, HDL, LDL, TG;
5. **Cardio-vascular**: SBP, DBP, HT, CHD, CP and MCP.

### 2.3. DNAm data quality control (QC) and preprocessing

In preprocessing raw DNAm data we mostly followed the procedures from [24] which are in line with the manufacturer’s recommended steps. In brief, we checked the array control probes’ metrics (Illumina Bead Control Reporter), signal detection *p*-values, and bead count numbers for all available cytosine-phospate-guanine (CpG) probes. Furthermore, we compared actual and DNAm predicted sex data for each sample. The samples included in this analysis were those where less than 1% of CpGs had a detection *p* ≥ 0.01, as well as having probes with a bead count number ≥ 3 and detection *p* < 0.01 in at least 99% of samples Initial DNAm data processing and QC data filtering were implemented using `R` v.4.1.0 [25] together with specialised `R` libraries `minfi` [26], `ChAMP` [27], and `ENmix` [28].

### 2.4. Epigenetic Age Acceleration

Epigenetic age acceleration (EAA) scores were calculated using the [DNA Methylation Online Calculator](https://dnamage.genetics.ucla.edu/) [3]. This web based tool gives nine EAAs based on five epigenetic scores, namely Horvath’s [3], Hannum’s [2], Skin and Blood [29], PhenoAge [4], and GrimAge [5] measures, see Table 1. Further details regarding epigenetic clocks and various EAAs are given in Appendix B.

**Table 1.** Summary of the EAA scores measured using DNA Methylation Online Calculator. **Abbreviations: CA** - chronological age, **EA** - epigenetic age, **EAA** - epigenetic age acceleration, **IEAA** - intrinsic epigenetic age acceleration, **EEAA** - extrinsic epigenetic age acceleration

| EAA | Clock | Info |
| --- | --- | --- |
| HannumAA | Hannum [2] | Residuals from regressing EA on CA |
| HannumEEAA | Hannum [30] | Residual from regressing the weighted average of Hannum's EA and estimated measures of blood cells counts on CA |
| HannumIEAA | Hannum [30] | Residuals from regressing Hannum's EA on CA and various blood immune cell counts |
| HorvathAAd | Horvath [3] | Difference between EA and CA |
| HorvathAAr | Horvath [3] | Residuals from regressing EA on CA |
| HorvathIEAA | Horvath [30] | Residuals from regressing Horvath's EA on CA and various blood immune cell counts |
| SkinBloodAA | Skin and Blood [29] | Residuals from regressing EA on CA |
| PhenoAA | PhenoAge [4] | Residuals from regressing EA on CA |
| GrimAA | GrimAge [5] | Residuals from regressing EA on CA |

### 2.5. Grouping

In this study we evaluated CVD-related phenotypes and their association with different EAA scores. It is expected that these phenotypes show small effect size (in comparison with some types of cancer) in blood DNA methylation, and hence in EAAs as well. Taking into account the relatively small sample size of our study, we decided to limit our analyses to the grouping of EAAs as described below.

Analysis of associations between EAAs and phenotypes in our study involves comparing the distributions of the phenotypic data in two groups. The grouping is based on binary split with respect to the sign of EAA, defined as follows:

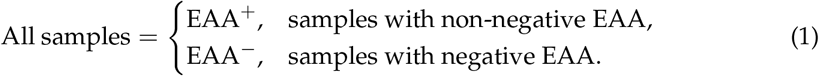

In other words, for each clock we use the definition (1) to split our cohort into two groups, one with EAA < 0 and the other with EAA ≥ 0, and then study the differences in phenotypic distribution between these groups. Similar grouping was also featured in previous studies based on EAAs [31,32].

### 2.6. Statistical Analysis

All statistical analyses were performed using `R` v.4.1.2. They include descriptive analysis of the available data using relevant techniques, such as univariate analysis, cross-tabulation, statistical hypothesis testing (Welch’s *t*-test [33] for continuous variables and Fisher’s exact test [34] for binary data), and linear regression-based data adjustments. Welch’s *t*-test null-hypothesis: mean values of a given variable in *EAA*^+^ and *EAA*^−^ groups are not different. Fisher’s exact test null-hypothesis: classifications of a given binary variable in EAA^+^ and EAA^−^ groups are not different. The significance level is defined as *α* = 0.05 for each EAA-phenotype association hypothesis test.

In order to consider the association between different EAAs and groups of phenotypes (all apart from the Lifestyle group), we controlled for family-wise error rate (FWER) using the Bonferroni correction [35,36], which was performed per group of phenotypes per EAA. The significance threshold for the Anthropometric, Metabolic, Lipids, and Cardio-vascular groups of phenotypes were calculated to be 0.025, 0.0166, 0.0125, and 0.0083, respectively. In the Lifestyle group we considered the smoking and alcohol intake data separately, thus, FWER-controlled significance threshold for alcohol consumption phenotypes is 0.025, and 0.05 for smoking status. It means that for each clock the group association was inferred from the individual phenotypes by controlling for FWER in the different phenotype groups. In other words, we define the EAA score to be associated with the group of phenotypes if for at least one of the phenotypes in the group the significance of the relationship is sustained with the Bonferroni-corrected threshold.

All the graphs presented in the paper were produced using `ggplot2` [37] and its extensions, `pheatmap` [38], `PerformanceAnalytics` [39] and base `R` functions.

## 3. Results

### 3.1. Associations between sex and phenotypes

Our dataset consists of (*n* = 306) samples (166 females and 140 males). Summaries of the dataset characteristics for all samples and for sex-specific groups are given in Tables A1 and A3. Table A1 contains descriptive statistics (range, mean, and standard deviation) for the available continuous phenotype data and the corresponding Welch’s *t*-test *p*-values and 95% confidence intervals. Table A3 includes count numbers and percentages for dichotomous variables, together with sex-specific odds ratios, 95% confidence intervals, and *p*-values calculated by performing a Fisher’s exact statistical test.

The Russian sample being considered in this study showed no significant difference between males and females in the distribution of chronological age, blood pressure values (both SBP and DBP), incidence of acute CHD events, diagnosis of hypertension and diabetes, or levels of triglycerides and fasting glucose. The phenotypes which were significantly different in males vs. females are anthropometric measures (BMI and WHR), lifestyle choices (alcohol consumption and smoking status), and blood levels of gamma-glutamyl transferase (GGT) and lipids (both LDL and HDL). Interestingly, in this Russian dataset there was no significant difference between the male and female odds ratios of being diagnosed with a carotid plaque (CP), but the odds ratios of having multiple carotid plaques (MCP) significantly differed between sexes.

### 3.2. EAAs are associated with some phenotypes and have strong sex bias

Our EAA analyses are based on nine EAA scores (described in Section 2.4) which were obtained from the five different epigenetic clock models, with multiple EAA scores derived from Horvath’s multi-tissue and Hannum’s clocks (3 EAAs each). Correlation coefficients are higher among EAAs based on the same clock than among EAAs from different clocks. Namely, Pearson correlation coefficients range between 0.78 and 0.97 within EAAs derived from Horvath’s and Hannum’s models, whilst the highest value of correlation for EAAs derived from separate clocks is *r* = 0.57, see correlation table on Figure A2. Note that neither of the EAAs is significantly correlated with chronological age apart from HorvathAAd, which is the only measure calculated without chronological age adjustment.

To explore connections among the variables we calculated correlation coefficients (Spearman correlation) and normalised entropy-based mutual information values for all the phenotypes and EAAs. Heatmaps for correlations (absolute values) and mutual information values, as well as correlations-based network plots are presented in corresponding Figures 1, A1, and 2, respectively. In both the correlation and mutual information plots the EAAs are clustered together with the exception of GrimAA, which displays very strong associations with sex and smoking status.

**Figure 1.**
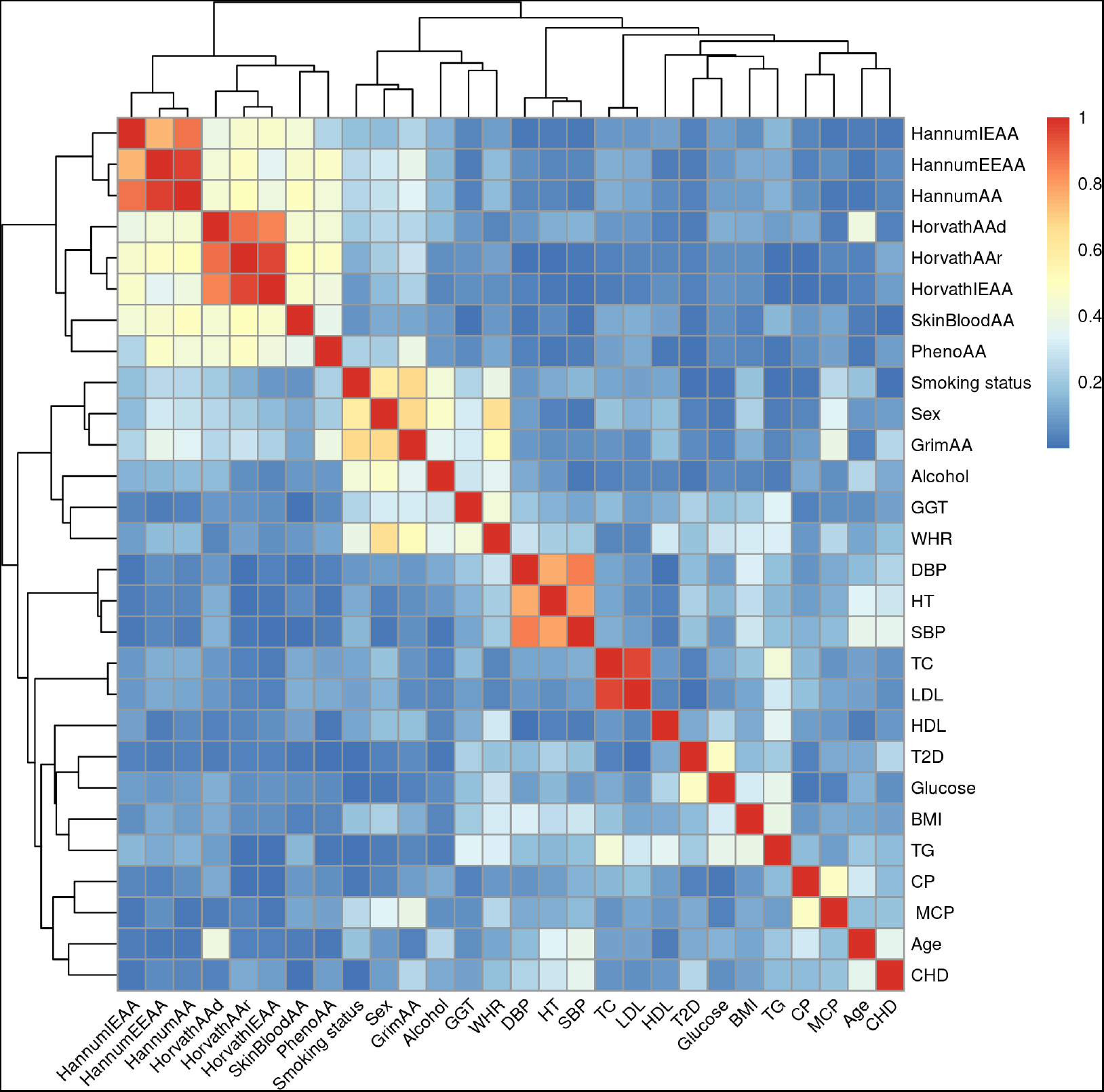
Heatmap of the correlations between all available traits and epigenetic age accelerations

**Figure 2.**
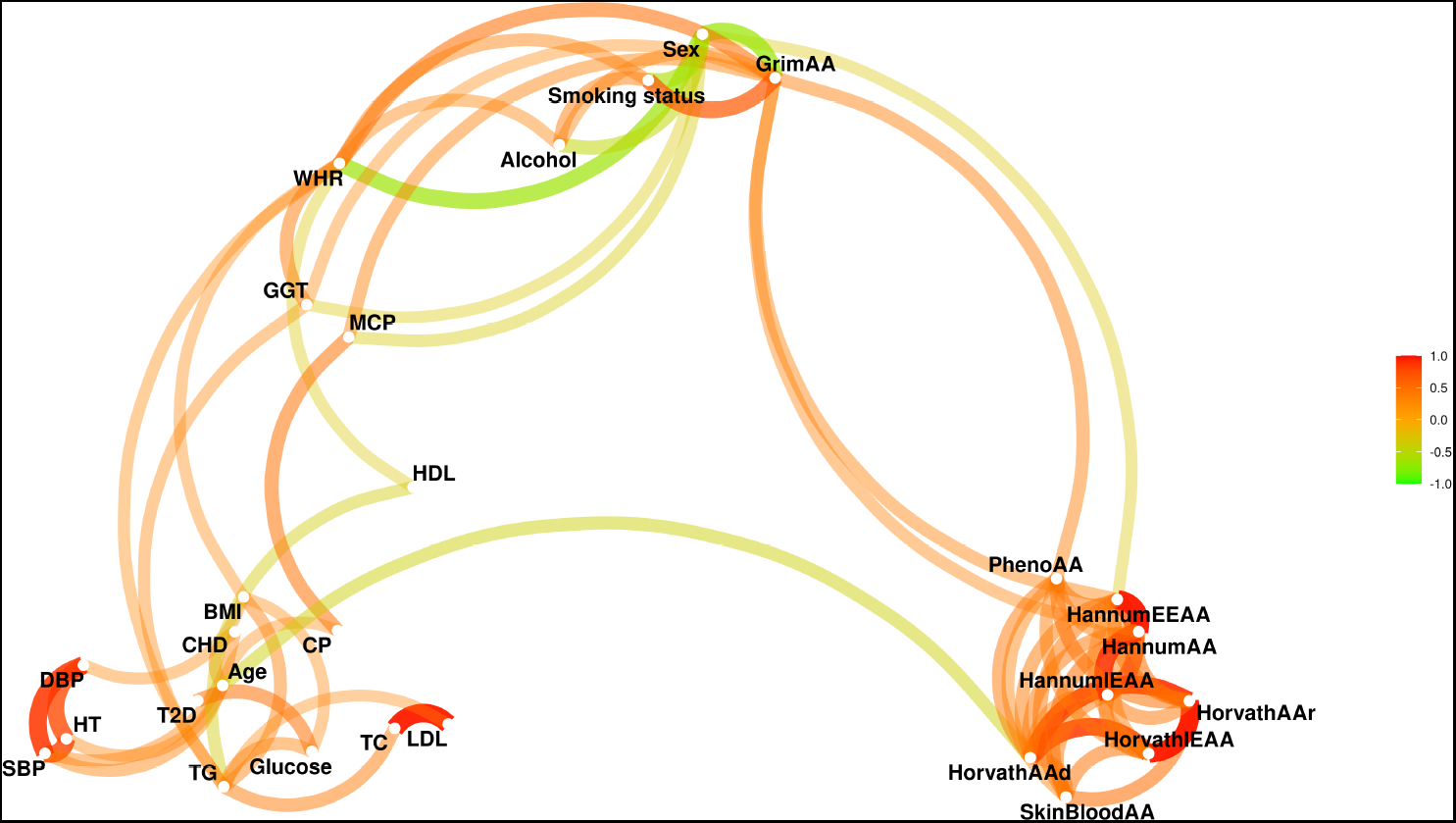
Network plot of of the connections among the phenotypes and EAAs in the dataset. Based on Spearman correlation coefficients with absolute values above 0.3

We further investigated the relationship between phenotypes and EAAs by splitting the dataset into EAA^+^ and EAA^−^ groups using (1), and, subsequently, testing the phenotype data distribution using a *t*-test for continuous variables, and Fisher’s exact test for binary variables. The corresponding statistical testing results are presented in Tables A5 and A6.

We noted that in nearly all EAA measures the size distribution of the EAA^+^ and EAA^−^ groups were within a 45%-55% range. The only exceptions to this were HorvathAAd (32% EAA^+^ samples vs. 68% EAA^−^ samples) and GrimAA (61% EAA^+^ samples vs. 39% EAA^−^ samples). Sex specific group splitting was found to be very unbalanced for all the EAAs for both sexes with an exception of HorvathIEAA, see Table A4. Furthermore, we observed significant differences in distributions of all nine EAA measures in our data between males and females, the corresponding data along with descriptive statistics are presented in Table A2. Given the strong association between sex and the various phenotypes examined in this study, the significant results obtained for EAA-phenotype associations might have been confounded by sex.

### 3.3. Sex-adjusted EAAs are associated with various phenotypes

In order to eliminate the undesired sex bias we adjusted all of the EAA scores by sex and then repeated the analyses described in the previous section, based on the calculated adjusted EAAs (adjEAA). Splitting the data into EAA^+^ and EAA^−^ resulted in balanced group sizes for all the adjEAAs, all of the groupings were within a 44%-56% range, see Table A4. The medians of the adjEAAs for sex-specific subsets located closer to 0 compared to the medians of unadjusted EAAs, see Figure 3**A**.

**Figure 3.**
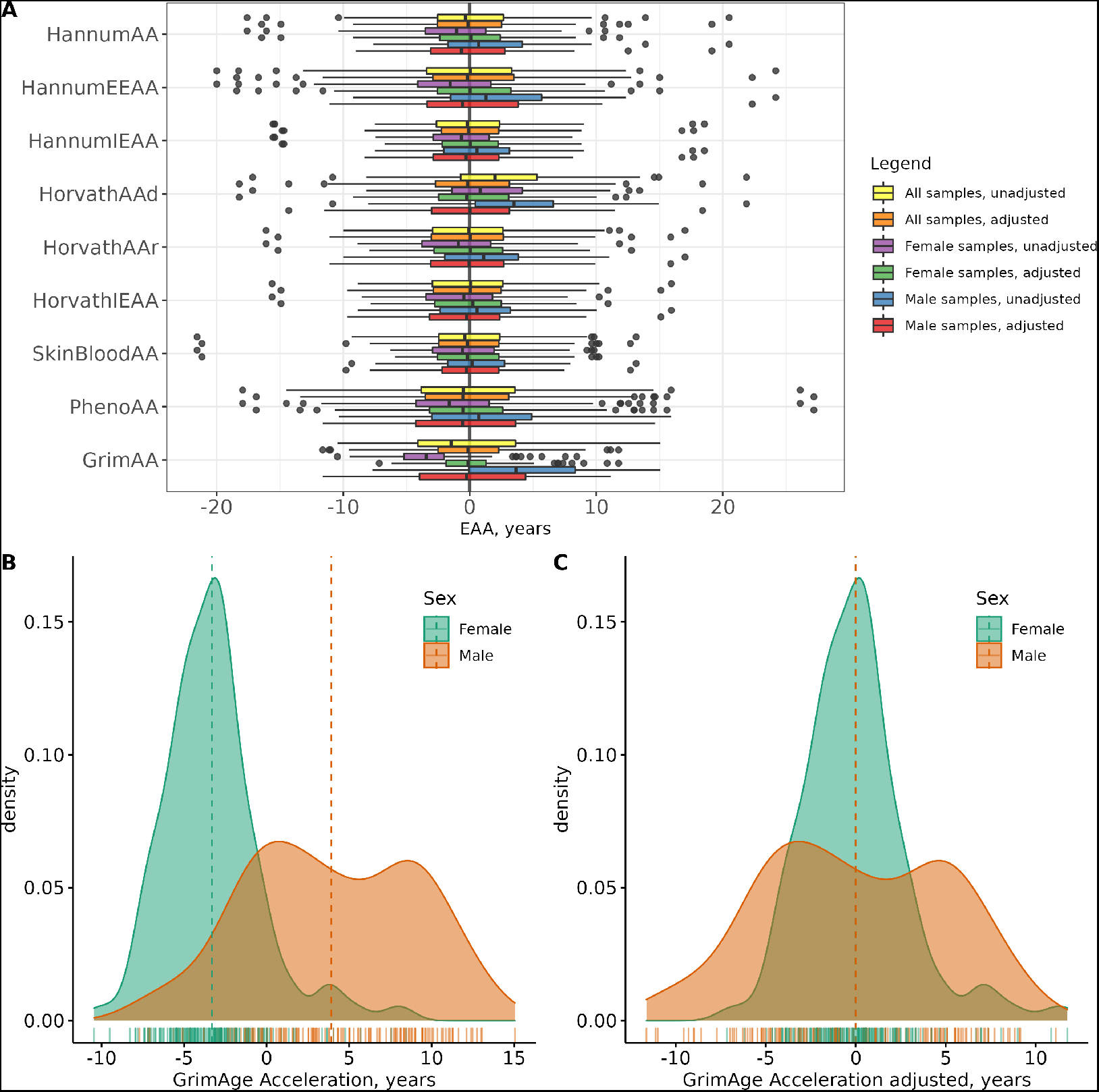
Boxplots of EAAs and sex-adjusted EAAs for all samples and sex-specific subsets (panel **A**), distribution of GrimAA distribution stratified by sex (panel **B**) and adjusted for sex (panel **C**).

The significant results from testing the differences in phenotype distribution between the EAA^+^ and EAA^−^ groups are given in Table 2. This table contains 95% confidence intervals, which indicate the trends in the direction of differences. The corresponding means and odds ratio values could be found in Table A6, where we present all testing outcomes regardless of their significance. For all available samples only four adjEAAs (GrimAA, PhenoAA, Horvath’s residuals, and IEAA) demonstrated statistically significant results for six phenotypes, with 4 phenotypes highlighted by GrimAA, and one phenotype each by the rest of the adjAAs (7 phenotype-EAA combinations in total). Among the differently distributed phenotypes are blood levels of GGT and LDL, smoking status and annual alcohol consumption, diagnosed MCP and incident CHD status; with the latter being the only phenotype that tested significantly different by multiple adjEAAs (GrimAA and Horvath’s differences). Interestingly, for the GrimAA clock, incident CHD and smoking status stayed significantly different for both male-only and female-only subsets, whilst GGT was not significantly different in any sex-specific groupings.

**Table 2.** Significant differences between groups with positive and negative epigenetic age acceleration scores (EAA^+^ and EAA^−^). Results which remained significant after controlling for family-wise error rate of 0.05 in groups of phenotypes per clock (as described in the Methods section 2.6) are presented in **bold**.

| Phenotype | EAA | *p*, all | 95% CI, all | *p*, F | 95% CI, F | *p*, M | 95% CI, M |
| --- | --- | --- | --- | --- | --- | --- | --- |
| *Anthropometric* |  |  |  |  |  |  |  |
| BMI | GrimAA |  |  |  |  | 0.039 | (-3.079, -0.079) |
| WHR | HorvathAAd |  |  | **0.004** | (-0.048, -0.009) |  |  |
|  | GrimAA |  |  | **0.010** | (0.006, 0.046) |  |  |
| *Lifestyle* |  |  |  |  |  |  |  |
| Smoking status | GrimAA | **<0.001** | (1.799, 4.895) | **0.026** | (1.077, 8.931) | **<0.001** | (4.5, 58.7) |
|  | HorvathAAd |  |  | **0.016** | (1.140, 9.454) |  |  |
|  | PhenoAA |  |  |  |  | **0.004** | (1.360, 8.319) |
| Alcohol, annual intake | HorvathIEAA | 0.028 | (-2832, -163) |  |  | **0.023** | (-5522, -422) |
|  | GrimAA |  |  |  |  | 0.049 | (16, 5370) |
| *Metabolic* |  |  |  |  |  |  |  |
| GGT | GrimAA | 0.023 | (0.728, 9.699) |  |  |  |  |
|  | HorvathAAr |  |  |  |  | 0.030 | (0.738, 14.5) |
| *Lipids* |  |  |  |  |  |  |  |
| TC | HannumAA |  |  | **0.009** | (-0.947, -0.141) |  |  |
|  | GrimAA |  |  | 0.046 | (0.008, 0.818) |  |  |
|  | PhenoAA |  |  | **0.010** | (-0.919, -0.127) |  |  |
|  | HannumEEAA |  |  | **0.003** | (-1.004, -0.203) |  |  |
| TG | GrimAA |  |  | 0.015 | (0.070, 0.632) |  |  |
| HDL | HorvathAAr |  |  | 0.013 | (0.026, 0.219) |  |  |
|  | SkinBloodAA |  |  | 0.027 | (0.012, 0.205) |  |  |
| LDL | PhenoAA | 0.037 | (-0.523, -0.016) | **0.004** | (-0.840, -0.157) |  |  |
|  | HannumAA |  |  | **0.010** | (-0.811, -0.112) |  |  |
|  | HannumEEAA |  |  | **0.002** | (-0.904, -0.215) |  |  |
| *Cardio-vascular* |  |  |  |  |  |  |  |
| CHD | GrimAA | **<0.001** | (1.518, 4.060) | **0.001** | (1.458, 5.955) | 0.042 | (1.020, 4.389) |
|  | HorvathAAr | **0.006** | (1.187, 3.139) |  |  | 0.018 | (1.150, 4.995) |
| CP | GrimAA |  |  |  |  | 0.009 | (1.367, 22.723) |
| MCP | GrimAA | **0.004** | (1.584, 26.779) |  |  | 0.009 | (1.401, 33.864) |
| HT | HorvathAAd |  |  | **0.005** | (0.202, 0.781) |  |  |
|  | GrimAA |  |  | 0.043 | (0.987, 3.764) |  |  |
| SBP | HorvathAAd |  |  | **0.008** | (-20.1, -3.2) |  |  |
|  | GrimAA |  |  | 0.024 | (1.3, 18.4) |  |  |
| DBP | HorvathAAd |  |  | **0.003** | (-10.626, -2.144) |  |  |
|  | GrimAA |  |  | 0.039 | (0.228, 8.832) |  |  |

Seven phenotype-EAA combinations were demonstrated to be statistically significant. Of these, five remained significant in male-only data subsets, and three remained significant in female-only subsets. In males the significant differences between the EAA^+^ and EAA^−^ groups were confirmed by four EAAs (the same for all samples) and seven phenotypes (10 phenotype-EAA combinations). Significant results for females feature seven EAAs (all apart from Horvath’s and HannumIEAA) and 10 traits (21 phenotype-EAA combinations). Nearly half (10 out of 21) of the results for the female subgroup presented in Table 2 relate to blood lipids measures (TG, LDL and HDL), and another 6 results for females relate to the presence of a hypertension diagnosis and blood pressure values (SBP and DBP). Neither lipidnor blood pressure-related phenotypes were associated with the EAA^+^/EAA^−^ grouping in males, unlike the presence of a CP/MCP diagnosis. Anthropometric parameters were also found to be statistically different in both sex-specific groups (BMI in males and WHR in females), but not for the combined dataset.

Our findings also suggest that some of the clocks are associated with groups of phenotypes. In particular, we observed that GrimAA and HorvathAAr are significantly associated with the cardio-vascular group of phenotypes for the entire dataset. In the female-only subset, HorvathAAd and GrimAA are both associated with the anthropometric and cardio-vascular groups, whilst PhenoAA, HannumAA, and HannumEEAA are associated with the lipids group. No significant results were found for the metabolic group or in the male-only subset.

### 3.4. Directions of some EAA-phenotype associations in sex-specific subsets are different

The results presented in Tables 2 and 3 summarise the significant outcomes of statistical hypothesis testing of our data based on grouping (1). Some of the phenotypes in the sex-specific groupings were highlighted by multiple EAAs, e.g. female total cholesterol (HannumAA, PhenoAA, GrimAA and EEAA) and male alcohol consumption (IEAA and GrimAA), however, the signs of the groups’ mean differences are not consistent.

**Table 3.**
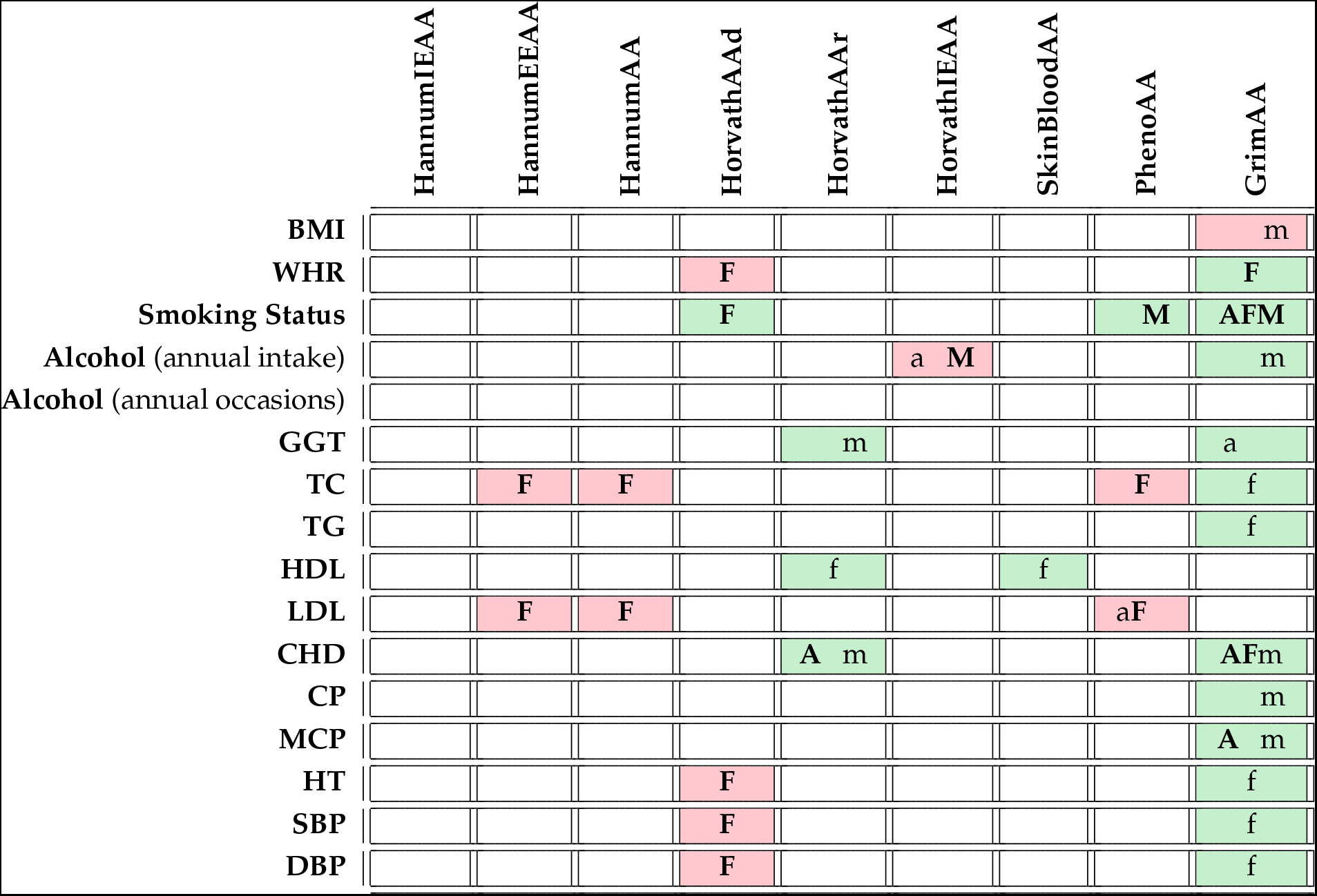
EAA-Phenotype association table. White colour indicates no significant association, green/red colour indicate significantly higher/lower values of phenotype measures (higher odds ratios) in *EAA*^+^ group compared to *EAA*^−^ for continuous (binary) phenotypes. Lower case letters indicate individual significant associations between a clock and a phenotype for all cohort participants (a), females (f), and males (m); capital letters indicate phenotype group significant association (controlled for family-wise error rate) for all (**A**), females (**F**), and males (**M**).

For instance, in Table 2, the confidence intervals for female WHR are positive for GrimAA and negative for HorvathAAd. Further investigation revealed that the EAA^+^ group have a higher mean WHR than in EAA^−^ for GrimAA, but the opposite is true for HorvathAAd, see Figure A6. Furthermore, the mean WHR values were higher in the EAA^−^ group for all three EAAs derived from Horvath’s clock, together with HannumIEAA and SkinBloodAA. Similar trends were observed in male annual alcohol consumption, see Figure A3, and in female levels of TC, HDL, LDL, and blood pressure values (SBP and DBP), see Figures A9, A8, A7, A12, and A11, respectively.

## 4. Discussion

The question of which EAA measure is the “best” or “most suitable” to study particular phenotypes is yet to be answered. For our data we decided to take into consideration all the EAA measures that could be calculated using [DNA Methylation Online Calculator](https://dnamage.genetics.ucla.edu/) [3], which is a relatively easy-to-use open-access tool. Epigenetic clocks included in the Online Calculator are featured in the vast majority of studies related to EAA-phenotype/disease associations (see e.g. [32,40–42]), as well as in benchmarking the newly developed DNAm-based clocks’ performance (see e.g.[43]).

Ability of the considered nine EAAs to reflect the differences in phenotype distribution was investigated by splitting the data based on the sign of the EAA scores. Similar grouping was also used in [31], where the risk of CHD was studied by splitting HorvathAAd and HannumAA to positive and negative groups. In recent paper [32], the authors used the positive/negative GrimAA and PhenoAA split to study the incident diabetes in the Coronary Artery Risk Development in Young Adults (CARDIA) cohort.

All the considered EAAs were independent (apart from HorvathAAd) of chronological age, but clearly sex-biased (Table A2), with generally lower EAA values for females. It was particularly obvious for the GrimAA scores, with distribution profiles separated for males and females, see Figure 3 **B-C**. As we pointed out in Section 3.3, splitting the dataset into EAA^+^ and EAA^−^ groups revealed big variation in group sizes for different scores (Table A4), which became particularly extreme for sex-specific subsets. To avoid unwanted confounding, for our analyses we adjusted EAAs by sex and proceeded with adjEAA values. This step resulted in more balanced EAA^+^/EAA^−^ group split for all adjEAAs. Of course, adjusting EAAs for sex did not affect the actual differences in phenotypes distributions between male and female subjects (see Tables A3 and A1). As a result, several phenotype-EAA combinations, which have previously demonstrated statistically significant results, did not persist after the adjustment (see Table A5).

Due to some phenotypes show sex-specific behaviour, see e.g. [44–46], we presented the results for males and females separately, alongside the results for the entire dataset. In one of the recent reviews [47], the authors pointed out the lack of sex-specific results involving EAAs and recommended splitting data by sex in downstream analyses. Our findings confirm the importance of using EAAs in sex-specific groups. We observed that the most phenotypes are reflected by some EAAs in one sex-specific group only. It should be noted that the results presented using positive and negative EAAs grouping (see Section 2.5). The observed association in groups could be further investigated on larger datasets using continuous EAAs.

We found that in our dataset both BMI and WHR were significantly different in males and females. Without adjusting for sex, multiple EAAs groupings highlighted significant differences in both BMI and WHR for all samples, but none of those associations replicated after adjustment (see Table A5). It is known [44], that in females WHR is associated with risks of CHD regardless of BMI, whilst in males WHR was found to be associated with incidence of CHD only for subjects with normal BMI measures. Our analysis found the anthropometric parameters to be statistically different in both sex-specific groups (BMI (GrimAA) in males and WHR (GrimAA and HorvathAA) difference in females), which is in line with results reported in large-scale US Sisters study [48] and Taiwan Biobank [49] cohorts. Interestingly (and opposite to findings in [44]), in [49] the authors report significant associations between WHR and EAAs (PhenoAge and GrimAge) in males, and between BMI and EAAs (PhenoAge and GrimAge) in females, which is the other way round in studied Russian sample (WHR in females and BMI in males). We would like to point out, that GrimAA grouping revealed higher mean WHR in female EAA^+^, but lower male mean BMI in the same group with positive EAAs (see Figures A6 and A5).

Lifestyle habits, including diet, smoking and alcohol consumption, are known to impact DNAm and being associated with epigenetic age in multiple studies, see e.g. [50–52]. Some DNAm clocks were specifically developed to be sensitive to smoking status, like, for example, GrimAge [5]. GrimAA was the only score associated with smoking status in the entire dataset, and the associations replicated in sex-specific subsets (Table 2). PhenoAA and HorvathAAd were also found to be significantly associated in male and female subgroups respectively. HorvathIEAA was significantly associated with annual alcohol consumption for all the samples and this association persisted in males, together with GrimAA, but not in females. Interestingly, the mean annual alcohol volume was higher in EAA^+^ group for GrimAA, but lower in EAA^+^ group for IEAA (Figure A3), which is not in line with the current state of the art in alcohol-ageing relationship, see e.g. review [53].

Previous publications suggest that EAAs are associated with diabetes and/or glucose levels [9,54,55]. It was also found that positive GrimAA (but not PhenoAA) is associated with higher 5-10 years incidence of type 2 diabetes, particularly for obese individuals [32]. In analysing this Russian sample we have not observed any significant associations of the considered EAAs with prevalent type 2 diabetes mellitus (T2DM) status and/or fasting blood glucose values. This might be attributed to the small proportion (11%) of the diabetics in our data compared to other studies (e.g. nearly 20% in [55]). Blood levels of GGT are associated with many dismetabolic conditions, including fatty liver, excessive alcohol consumption, increased risks of CHD and T2DM [56,57], and is known to be different in men and women [45], with no unified reference values. For the entire dataset, GrimAA EAA^+^/EAA^−^ grouping demonstrated significant difference in serum GGT measures. This result was not replicated in sex-specific subsets, but at the same time, in male subgroup GGT level difference was detected in HorvathAAr split (see Figure A4).

Blood levels of total cholesterol, TG and lipoproteins (HDL and LDL) are known to be sex-specific and associated with risk of developing CVD in both sexes, see e.g. [58]. Changes in lipids concentrations are also shown to be reflected in age-related changes in DNAm following dietary interventions [59]. Furthermore, associations of EAAs and lipids levels were confirmed in several studies [60,61]. In our entire dataset, among all available lipids data, only mean LDL levels in EAA^+^ with PhenoAA grouping were significantly lower than in EAA^−^, and this result persisted in female subset. No significant differences in mean lipids concentrations were highlighted by any EAA split for the male subgroup, whilst ten EAA-lipids phenotypes associations were highlighted in females. In particular, in female subset GrimAA grouping demonstrated significantly higher group mean levels of total cholesterol and TG in EAA^+^ compared to EAA^−^ (see Table 2, Figures A9 and A10). At the same time mean TC and LDL concentrations (Figure A7) were significantly lower in EAA^+^ group in PhenoAA, HannumAA and EEAA splits. Female HDL levels associations were picked up in SkinBloodAA and HorvathAAr groupings, with higher lipoprotein concentration in EAA^+^ group (see Figure A8). Remarkably, for all four considered lipids-related measures, known CVD risk factors (high TC, LDL, TG, and low HDL) were associated (not all significantly) with positive age acceleration only for GrimAA grouping, whilst the opposite was demonstrated in all the significant (and vast majority of insignificant) EAAs-lipids associations based on other EAA splits (see Figures A7, A8, A9 and A10). In view of recently published age-related sex-specific trends in lipid levels [62] and hypertension prevalence [46], it would be interesting to conduct an extended sex-specific analyses on EAA-lipids and hypertension associations for the particular age groups to see whether EAA values reflect the observed age-related patterns.

Data on carotid atherosclerosis and advanced atherosclerosis, which are defined in our study as the presence of at least a single (CP) and multiple carotid plaques (MCP) respectively, was available for only 34% of the participants, with 50/23/14 and 55/23/2 total/CP/MCP samples available for males and females. Only GrimAA grouping was significantly associated with CP in males and MCP in the entire dataset and its male only subset. In female specific subset blood pressure values (both SBP and DBP) and hypertension status were significantly associated with HorvathAAd and GrimAA groupings. Interestingly, in case of GrimAA group split, mean values of SBP and DBP were higher in EAA^+^ group, which might indicate the increased risk of CVD [63]. This is the opposite to the corresponding results of HorvathAAd grouping. None of these phenotypes were highlighted in the entire dataset and male subset. Two groupings, GrimAA and HorvathAAr, were significantly associated with incident CHD for all available samples. The results persisted in male subset for both groupings and in female subset for GrimAA split only. Similar results were also described in Genetic Epidemiology Network of Arteriopathy (GENOA) dataset study [55], where the authors reported significant connections not only between GrimAA and incident CVD, but also between GrimAA and time to the CVD event.

Notably, while higher odds of CHD were associated with EAA^+^ for both GrimAA and HorvathAAr, only GrimAA EAA^+^ was consistently associated with more harmful phenotypes values, indicating higher risk of CHD. All other EAA splits demonstrated mostly the opposite behaviour regarding available risk factors (lipids, anthropometric, lifestyle and cardio-vascular).

The observed associations of the various EAAs with both individual and groups of phenotypes suggest that epigenetic age acceleration scores are sensitive to various cardiometabolic parameters, which might indicate their prognostic potential for related disorders. Further investigations conducted on well-annotated larger datasets are needed to improve the understanding the mechanisms behind those associations, and, possibly developing new biomarkers. These might be extended by applying other epigenetic age models and using continuous EAAs in association studies.

## 5. Conclusions

Our study conducted on a subset of HAPIEE cohort shows that EAAs are sex-specific and should be adjusted for sex in EAA-phenotypes association studies. Moreover, even after adjusting for sex, the associations between EAAs and considered 18 cardiometabolic phenotypes are sex-specific. The only two phenotype-EAA associations persisted through the entire dataset and both male and female subsets are incident CHD and smoking status.

Among all considered epigenetic clocks, GrimAge was significantly associated with more phenotypes than any other EA scores, but for most of the phenotypes those associations are weaker than in other scores. Furthermore, for some EAAs, the direction of the association with phenotype is counter-intuitive, i.e. lower EAA scores corresponded to more harmful values of the phenotypes. The observed associations of the various EAAs with both individual and groups of phenotypes suggest that epigenetic age acceleration scores are sensitive to various cardio-metabolic parameters, which might indicate their prognostic potential for related disorders.

## Author Contributions

OC drafted the manuscript with the input from all the authors. EC, KP and TAW equally contributed to all aspects of data analysis and draft preparation. NH, JS, AR, VM, MB, HP, VV, SM, and SB participated in data generation, analyses design and manuscript preparation. VV, SM and SB supervised the study. All authors read and approved the manuscript.

## Funding

The baseline HAPIEE study was funded by the Welcome Trust (WT064947, WT081081, 106554/Z/14/Z), the US National Institute of Aging (1RO1AG23522). O. Chervova and S. Beck were supported by grants from the Frances and Augustus Newman Foundation (172074). O. Chervova, S. Beck, M. Bobak and H. Pikhart were supported by EU-H2020 Project “CETOCOEN Excellence” (857560). T.A. Widayati was funded by Indonesian Endowment Fund (Lembaga Pengelola Dana Pendidikan).

## Institutional Review Board Statement

The study was conducted in accordance with the Declaration of Helsinki, and approved by the Institutional Ethics Committee of IIPM—Branch of IC&G SB RAS (Institute of Internal and Preventive Medicine—Branch of Federal State Budgeted Research Institution, “Federal Research Center, Institute of Cytology and Genetics, Siberian Branch of the Russian Academy of Sciences”), Protocol no. 1 from 14 March 2002 and Protocol no. 12 from 8 December 2020.

## Informed Consent Statement

Informed consent was obtained from all subjects involved in the study.

## Data Availability Statement

Raw DNA methylation IDAT files have been deposited at the European Genome-phenome Archive (EGA), which is hosted by the EBI and the CRG, under accession number EGAS00001006390. The dataset is under embargo at the moment. Further information about EGA can be found on https://ega-archive.org “The European Genome-phenome Archive of human data consented for biomedical research” [64].

## Acknowledgments

The authors are grateful to UCL Cancer Institute Medical Genomics Lab members for stimulating discussions.

## Conflicts of Interest

The authors declare no conflict of interest. The funders had no role in the design of the study; in the collection, analyses, or interpretation of data; in the writing of the manuscript; or in the decision to publish the results.

## Abbreviations

The following abbreviations are used in this manuscript:

ACS: Acute coronary syndrome
adjEAA: Adjusted epigenetic age acceleration
ASE: American Society of Echocardiography
BMI: Body mass index
BP: Blood pressure
CA: Chronological age
CARDIA: Coronary Artery Risk Development in Young Adults
CHD: Coronary heart disease
CP: Carotid plaque
CpG: Cytosine-phospate-guanine
CVD: Cardio-vascular disease
DBP: Diastolic blood pressure
DNAm: DNA methylation
EA: Epigenetic age
EAA: Epigenetic age acceleration
EASD: European Association for the Study of Diabetes
EEAA: Extrinsic epigenetic age acceleration
EGA: European Genome-phenome Archive
ESC: European Society of Cardiology
ESH: European Society of Hypertension
FPG: Fasting plasma glucose
GENOA: Genetic Epidemiology Network of Arteriopathy
GGT: Gamma-glutamyl transferase
HAPIEE: Health, Alcohol, and Psychosocial Factors in Eastern Europe
HDL: High-density lipoprotein
HT: Hypertension
IEAA: Intrinsic epigenetic age acceleration
IIPM: Institute of Internal and Preventive Medicine
LDL: Low-density lipoprotein
MCP: Multiple carotid plaques
MI: Myocardial infarction
QC: Quality control
SBP: Systolic blood pressure
T2DM: Type 2 diabetes mellitus
TC: Total cholesterol
TG: Triglycerides
WHR: Waist-hip ratio

## Appendix A. Supplementary Figures and Tables

**Table A1.**
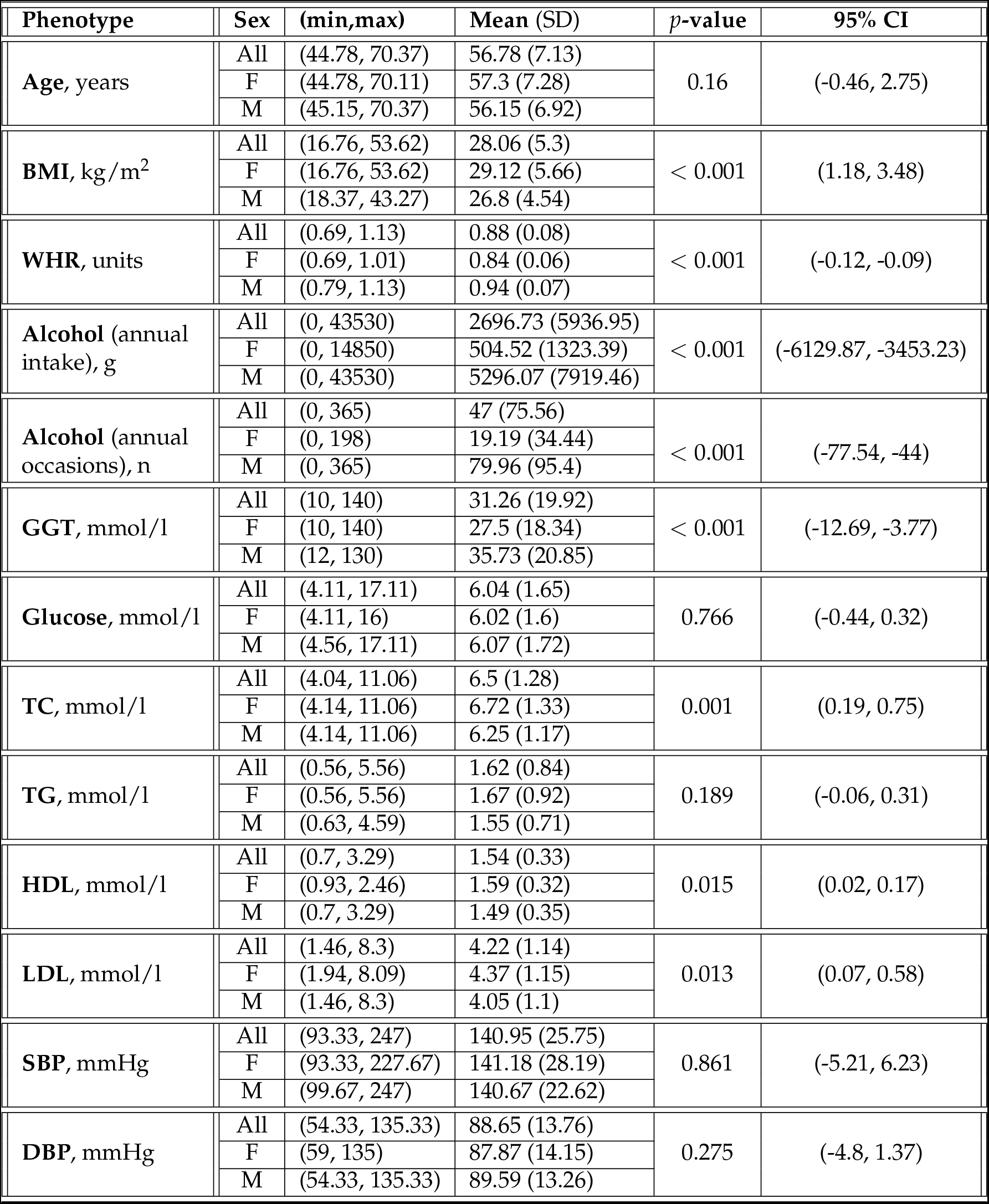
Summary of the continuous phenotype data for males and females. Sample size is *n* = 306, 166/140 female/male. Glucose measurements are only available for *n* = 298, 159/139 female/male samples. *p*-values obtained from the Welch’s *t*-test testing difference between male and female groups for each variable (**H**_0_: Mean value of the variable is the same for male and female groups).

| Phenotype | Sex | (min,max) | Mean (SD) | $p$-value | 95% CI |
| --- | --- | --- | --- | --- | --- |
| Age, years | All | (44.78, 70.37) | 56.78 (7.13) | 0.16 | (-0.46, 2.75) |
|  | F | (44.78, 70.11) | 57.3 (7.28) |  |  |
|  | M | (45.15, 70.37) | 56.15 (6.92) |  |  |
| BMI, kg/m<sup>2</sup> | All | (16.76, 53.62) | 28.06 (5.3) | < 0.001 | (1.18, 3.48) |
|  | F | (16.76, 53.62) | 29.12 (5.66) |  |  |
|  | M | (18.37, 43.27) | 26.8 (4.54) |  |  |
| WHR, units | All | (0.69, 1.13) | 0.88 (0.08) | < 0.001 | (-0.12, -0.09) |
|  | F | (0.69, 1.01) | 0.84 (0.06) |  |  |
|  | M | (0.79, 1.13) | 0.94 (0.07) |  |  |
| Alcohol (annual intake), g | All | (0, 43530) | 2696.73 (5936.95) | < 0.001 | (-6129.87, -3453.23) |
|  | F | (0, 14850) | 504.52 (1323.39) |  |  |
|  | M | (0, 43530) | 5296.07 (7919.46) |  |  |
| Alcohol (annual occasions), n | All | (0, 365) | 47 (75.56) | < 0.001 | (-77.54, -44) |
|  | F | (0, 198) | 19.19 (34.44) |  |  |
|  | M | (0, 365) | 79.96 (95.4) |  |  |
| GGT, mmol/l | All | (10, 140) | 31.26 (19.92) | < 0.001 | (-12.69, -3.77) |
|  | F | (10, 140) | 27.5 (18.34) |  |  |
|  | M | (12, 130) | 35.73 (20.85) |  |  |
| Glucose, mmol/l | All | (4.11, 17.11) | 6.04 (1.65) | 0.766 | (-0.44, 0.32) |
|  | F | (4.11, 16) | 6.02 (1.6) |  |  |
|  | M | (4.56, 17.11) | 6.07 (1.72) |  |  |
| TC, mmol/l | All | (4.04, 11.06) | 6.5 (1.28) | 0.001 | (0.19, 0.75) |
|  | F | (4.14, 11.06) | 6.72 (1.33) |  |  |
|  | M | (4.14, 11.06) | 6.25 (1.17) |  |  |
| TG, mmol/l | All | (0.56, 5.56) | 1.62 (0.84) | 0.189 | (-0.06, 0.31) |
|  | F | (0.56, 5.56) | 1.67 (0.92) |  |  |
|  | M | (0.63, 4.59) | 1.55 (0.71) |  |  |
| HDL, mmol/l | All | (0.7, 3.29) | 1.54 (0.33) | 0.015 | (0.02, 0.17) |
|  | F | (0.93, 2.46) | 1.59 (0.32) |  |  |
|  | M | (0.7, 3.29) | 1.49 (0.35) |  |  |
| LDL, mmol/l | All | (1.46, 8.3) | 4.22 (1.14) | 0.013 | (0.07, 0.58) |
|  | F | (1.94, 8.09) | 4.37 (1.15) |  |  |
|  | M | (1.46, 8.3) | 4.05 (1.1) |  |  |
| SBP, mmHg | All | (93.33, 247) | 140.95 (25.75) | 0.861 | (-5.21, 6.23) |
|  | F | (93.33, 227.67) | 141.18 (28.19) |  |  |
|  | M | (99.67, 247) | 140.67 (22.62) |  |  |
| DBP, mmHg | All | (54.33, 135.33) | 88.65 (13.76) | 0.275 | (-4.8, 1.37) |
|  | F | (59, 135) | 87.87 (14.15) |  |  |
|  | M | (54.33, 135.33) | 89.59 (13.26) |  |  |

**Table A2.**
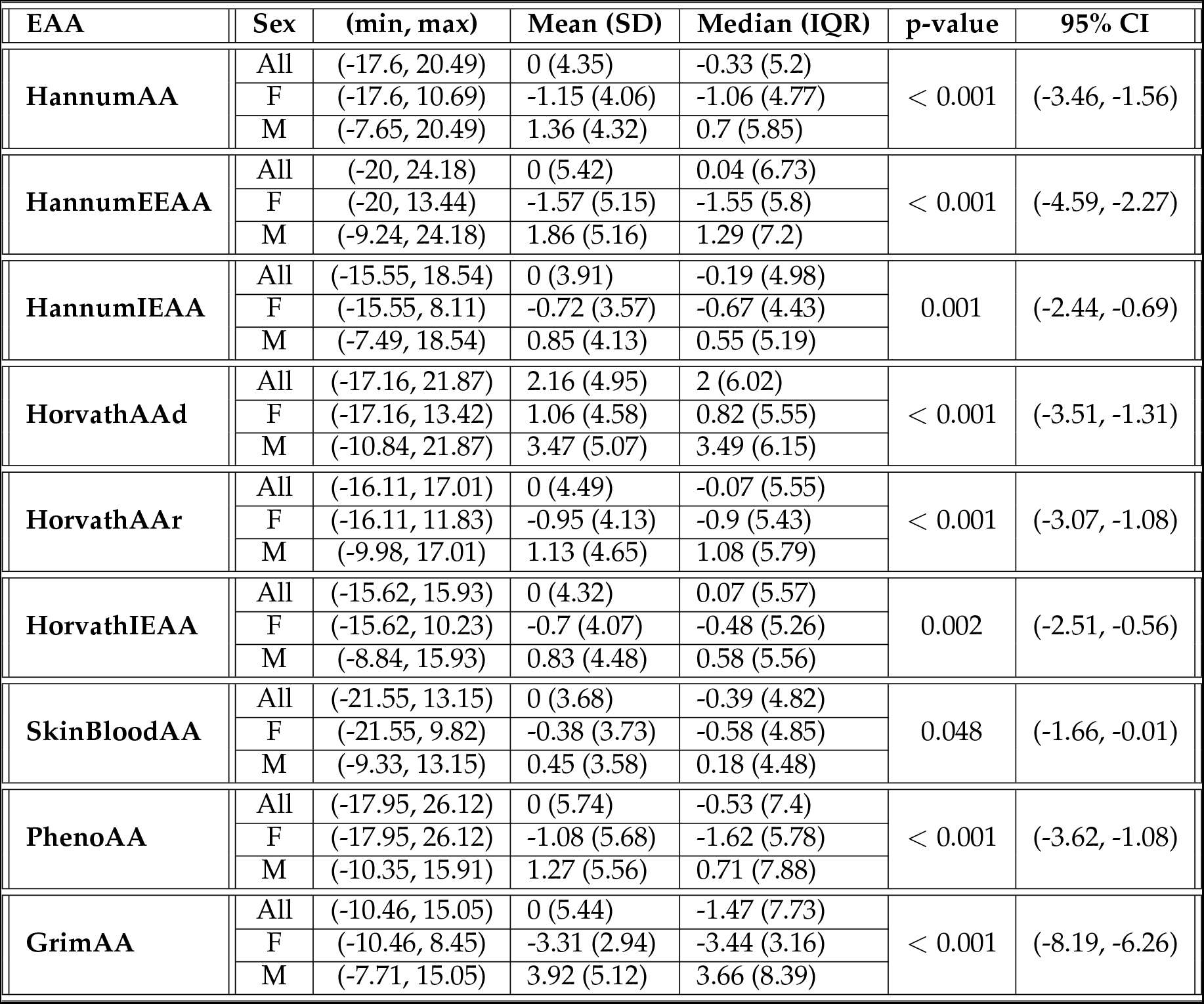
Summary of EAAs scores data for males and females. EAA scores before adjustment for sex. *p*-values obtained from the Welch’s *t*-test testing difference between male and female groups for each EAA (**H**_0_: Mean value of the EAA is the same for male and female groups).

| EAA | Sex | (min, max) | Mean (SD) | Median (IQR) | p-value | 95% CI |
| --- | --- | --- | --- | --- | --- | --- |
| **HannumAA** | All | (-17.6, 20.49) | 0 (4.35) | -0.33 (5.2) | < 0.001 | (-3.46, -1.56) |
|  | F | (-17.6, 10.69) | -1.15 (4.06) | -1.06 (4.77) |  |  |
|  | M | (-7.65, 20.49) | 1.36 (4.32) | 0.7 (5.85) |  |  |
| **HannumEEAA** | All | (-20, 24.18) | 0 (5.42) | 0.04 (6.73) | < 0.001 | (-4.59, -2.27) |
|  | F | (-20, 13.44) | -1.57 (5.15) | -1.55 (5.8) |  |  |
|  | M | (-9.24, 24.18) | 1.86 (5.16) | 1.29 (7.2) |  |  |
| **HannumIEAA** | All | (-15.55, 18.54) | 0 (3.91) | -0.19 (4.98) | 0.001 | (-2.44, -0.69) |
|  | F | (-15.55, 8.11) | -0.72 (3.57) | -0.67 (4.43) |  |  |
|  | M | (-7.49, 18.54) | 0.85 (4.13) | 0.55 (5.19) |  |  |
| **HorvathAAd** | All | (-17.16, 21.87) | 2.16 (4.95) | 2 (6.02) | < 0.001 | (-3.51, -1.31) |
|  | F | (-17.16, 13.42) | 1.06 (4.58) | 0.82 (5.55) |  |  |
|  | M | (-10.84, 21.87) | 3.47 (5.07) | 3.49 (6.15) |  |  |
| **HorvathAAr** | All | (-16.11, 17.01) | 0 (4.49) | -0.07 (5.55) | < 0.001 | (-3.07, -1.08) |
|  | F | (-16.11, 11.83) | -0.95 (4.13) | -0.9 (5.43) |  |  |
|  | M | (-9.98, 17.01) | 1.13 (4.65) | 1.08 (5.79) |  |  |
| **HorvathIEAA** | All | (-15.62, 15.93) | 0 (4.32) | 0.07 (5.57) | 0.002 | (-2.51, -0.56) |
|  | F | (-15.62, 10.23) | -0.7 (4.07) | -0.48 (5.26) |  |  |
|  | M | (-8.84, 15.93) | 0.83 (4.48) | 0.58 (5.56) |  |  |
| **SkinBloodAA** | All | (-21.55, 13.15) | 0 (3.68) | -0.39 (4.82) | 0.048 | (-1.66, -0.01) |
|  | F | (-21.55, 9.82) | -0.38 (3.73) | -0.58 (4.85) |  |  |
|  | M | (-9.33, 13.15) | 0.45 (3.58) | 0.18 (4.48) |  |  |
| **PhenoAA** | All | (-17.95, 26.12) | 0 (5.74) | -0.53 (7.4) | < 0.001 | (-3.62, -1.08) |
|  | F | (-17.95, 26.12) | -1.08 (5.68) | -1.62 (5.78) |  |  |
|  | M | (-10.35, 15.91) | 1.27 (5.56) | 0.71 (7.88) |  |  |
| **GrimAA** | All | (-10.46, 15.05) | 0 (5.44) | -1.47 (7.73) | < 0.001 | (-8.19, -6.26) |
|  | F | (-10.46, 8.45) | -3.31 (2.94) | -3.44 (3.16) |  |  |
|  | M | (-7.71, 15.05) | 3.92 (5.12) | 3.66 (8.39) |  |  |

**Figure A1.**
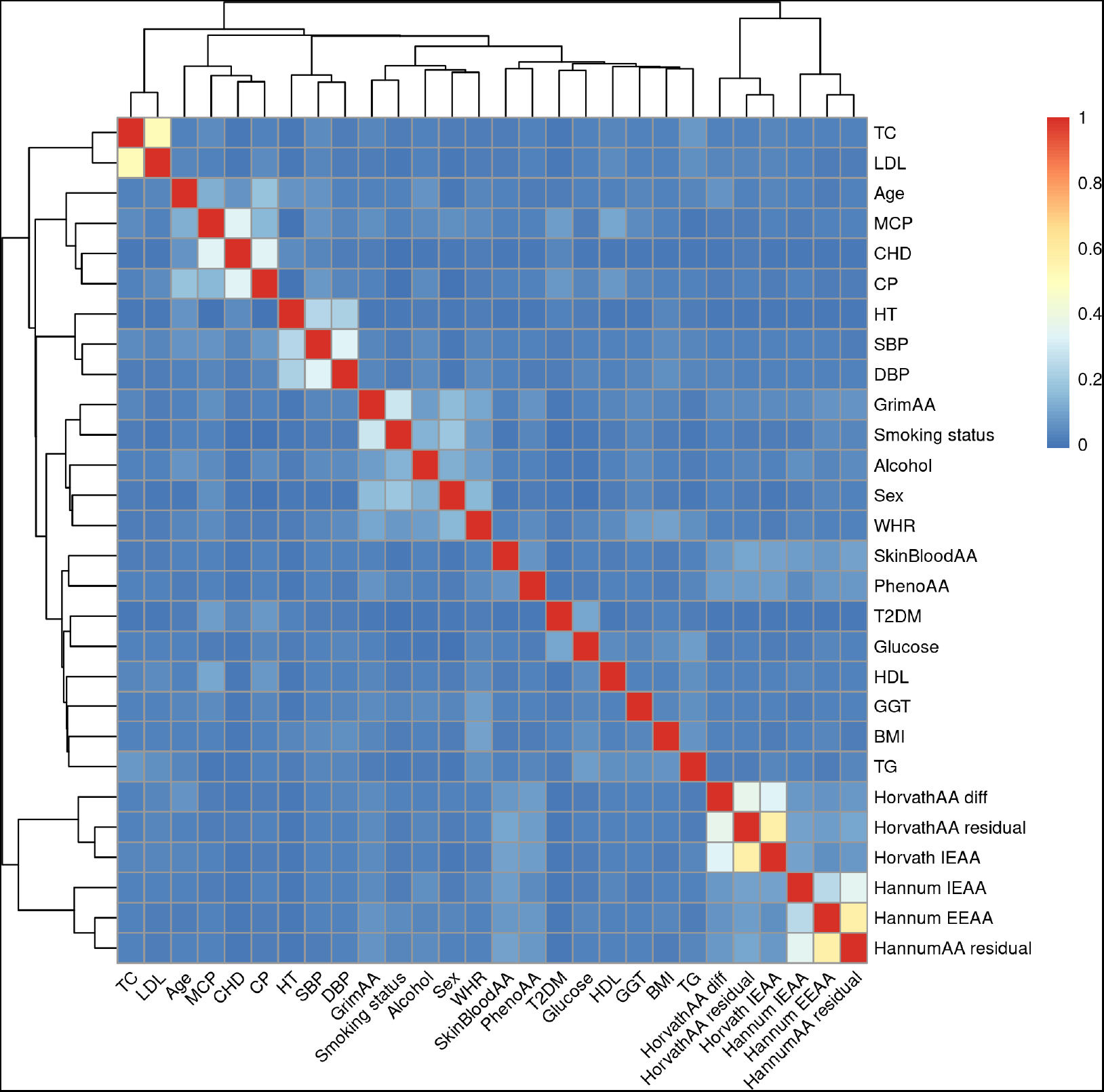
Normalised mutual information heatmap.

**Figure A2.**
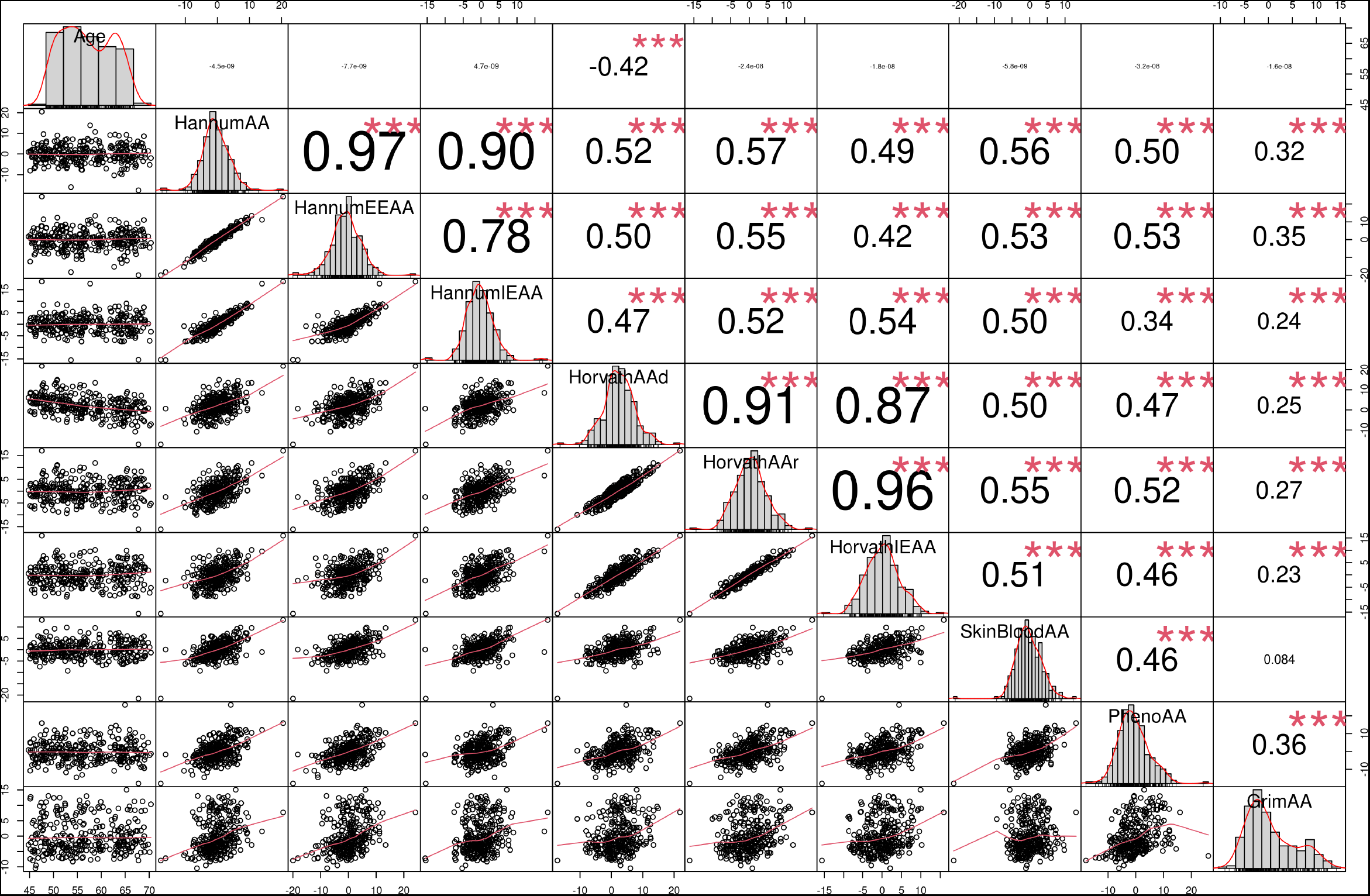
Correlation table for EAAs

**Figure A3.**
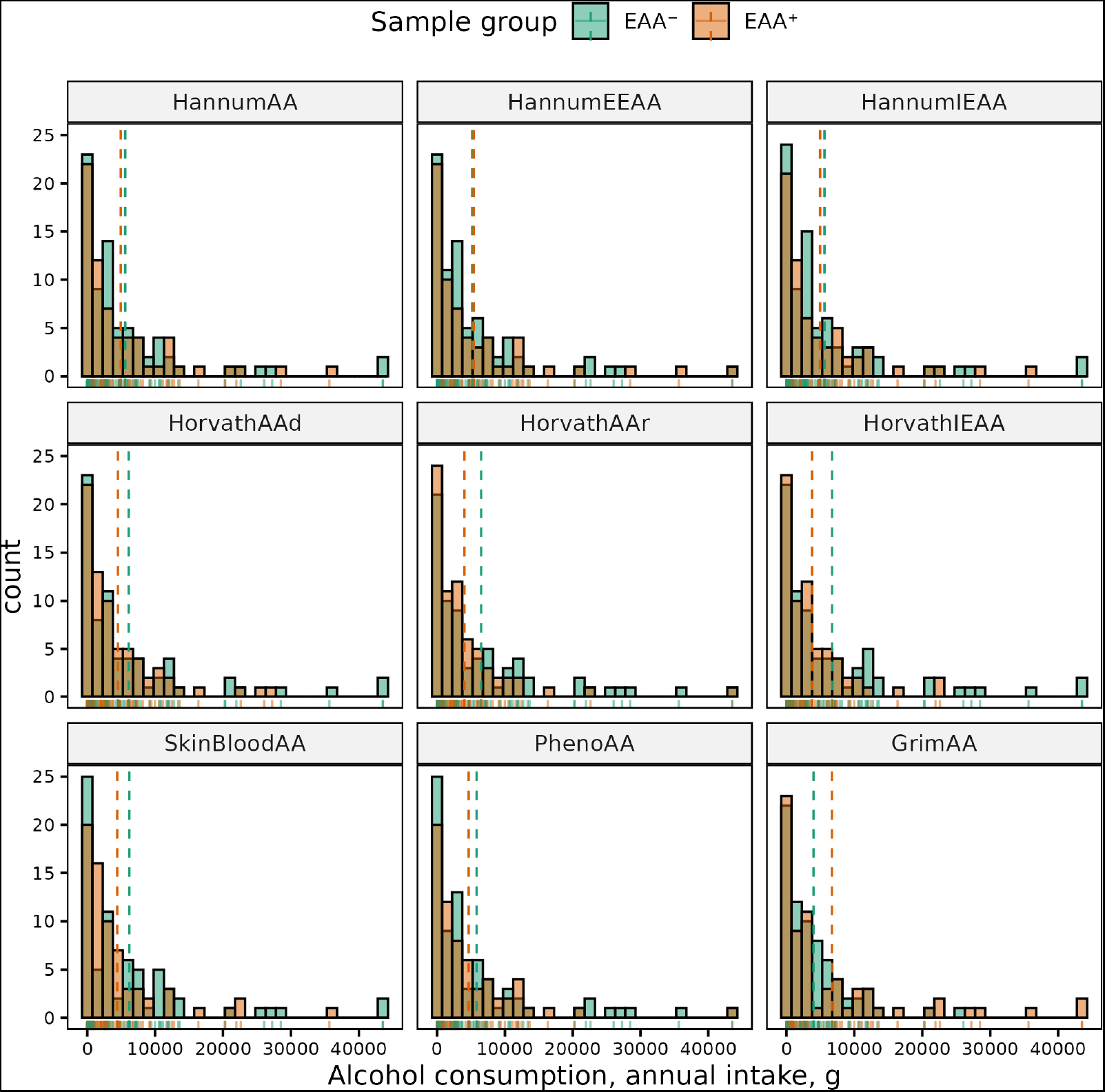
Histograms of distributions of annual alcohol consumption in males in EAA^+^ and EAA^−^ groups. Dashed lines correspond to the group means.

**Figure A4.**
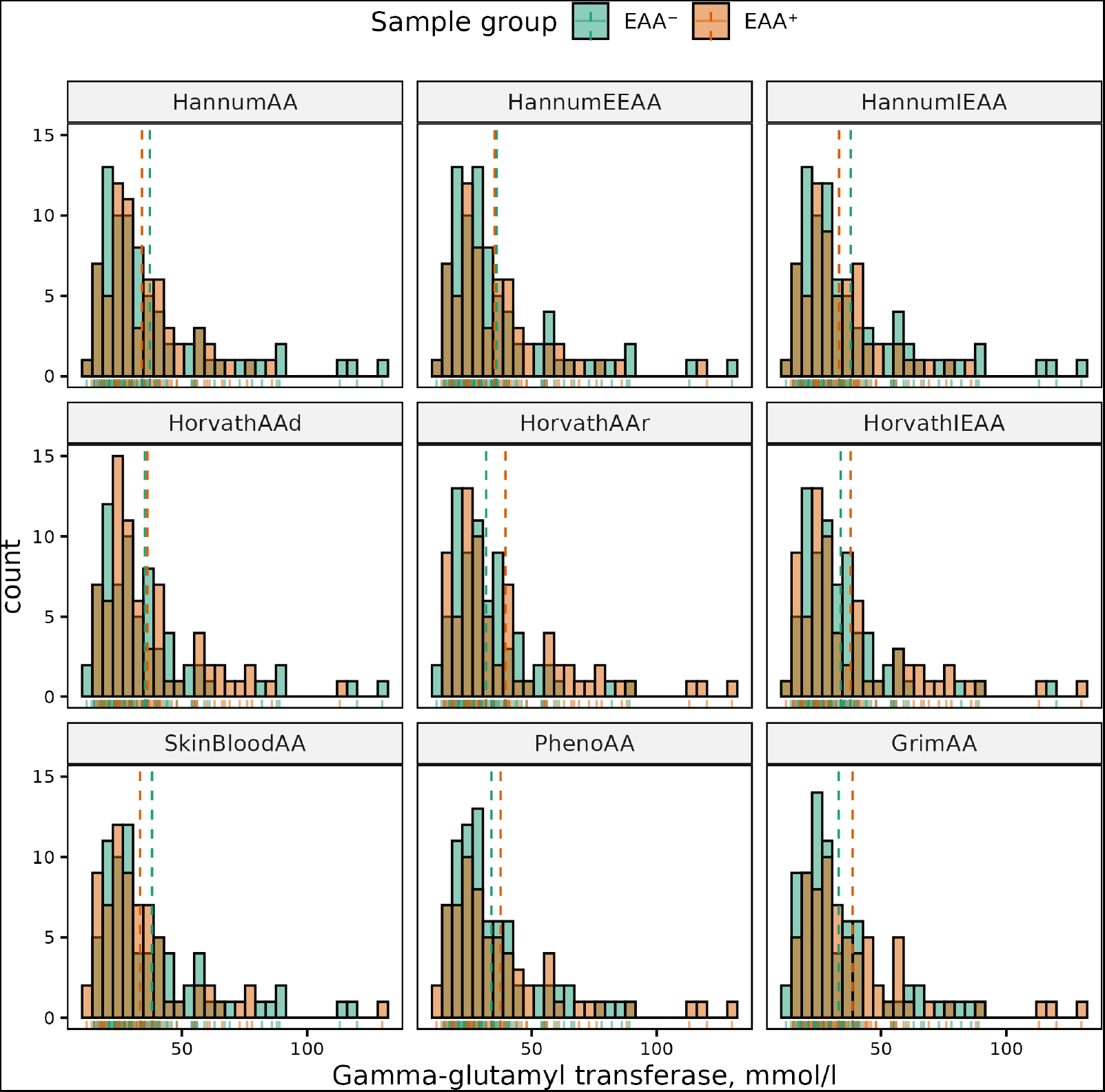
Histograms of distributions of GGT levels in males in EAA^+^ and EAA^−^ groups. Dashed lines correspond to the group means.

**Figure A5.**
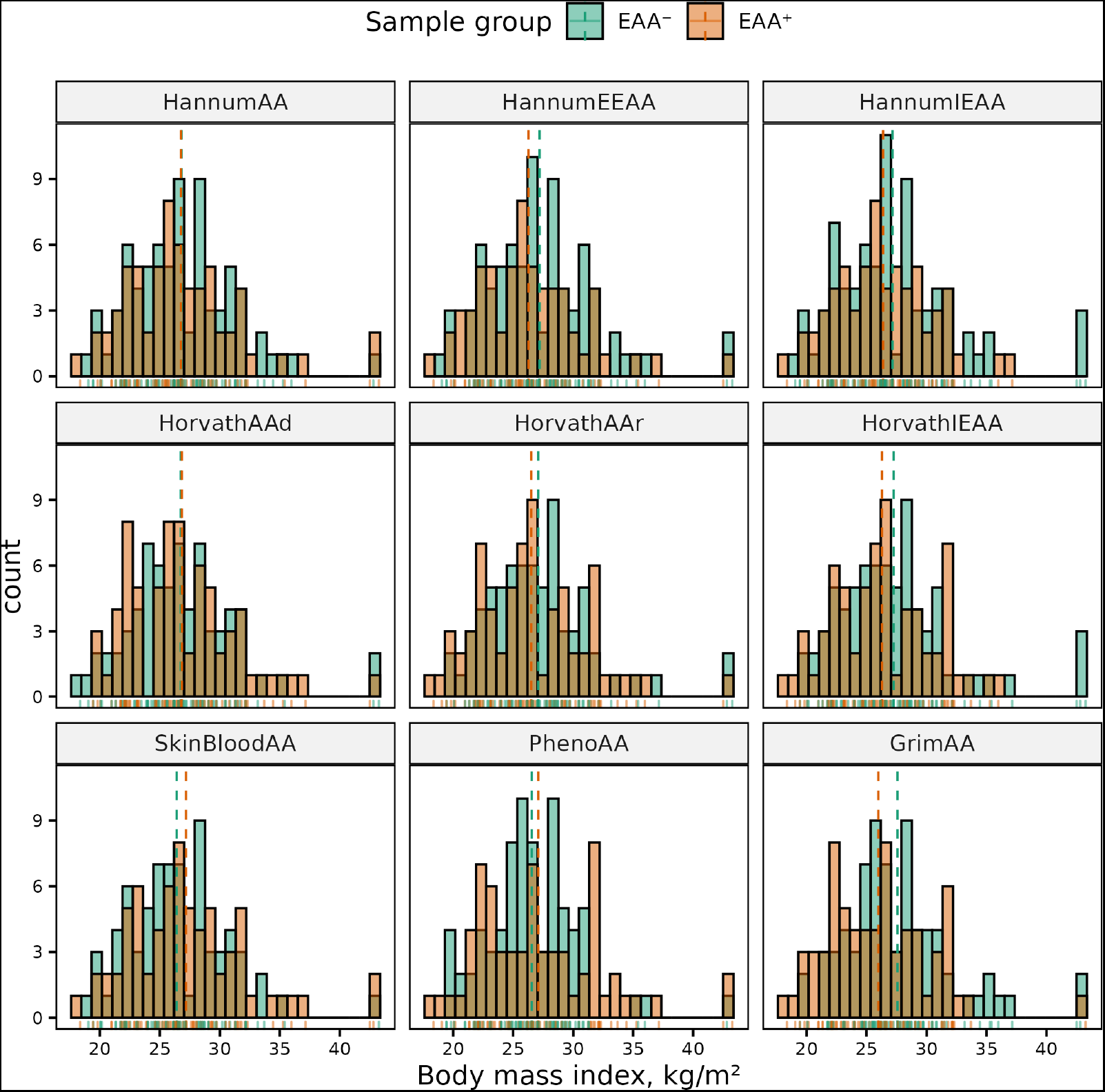
Histograms of distributions of BMI in males in EAA^+^ and EAA^−^ groups. Dashed lines correspond to the group means.

**Figure A6.**
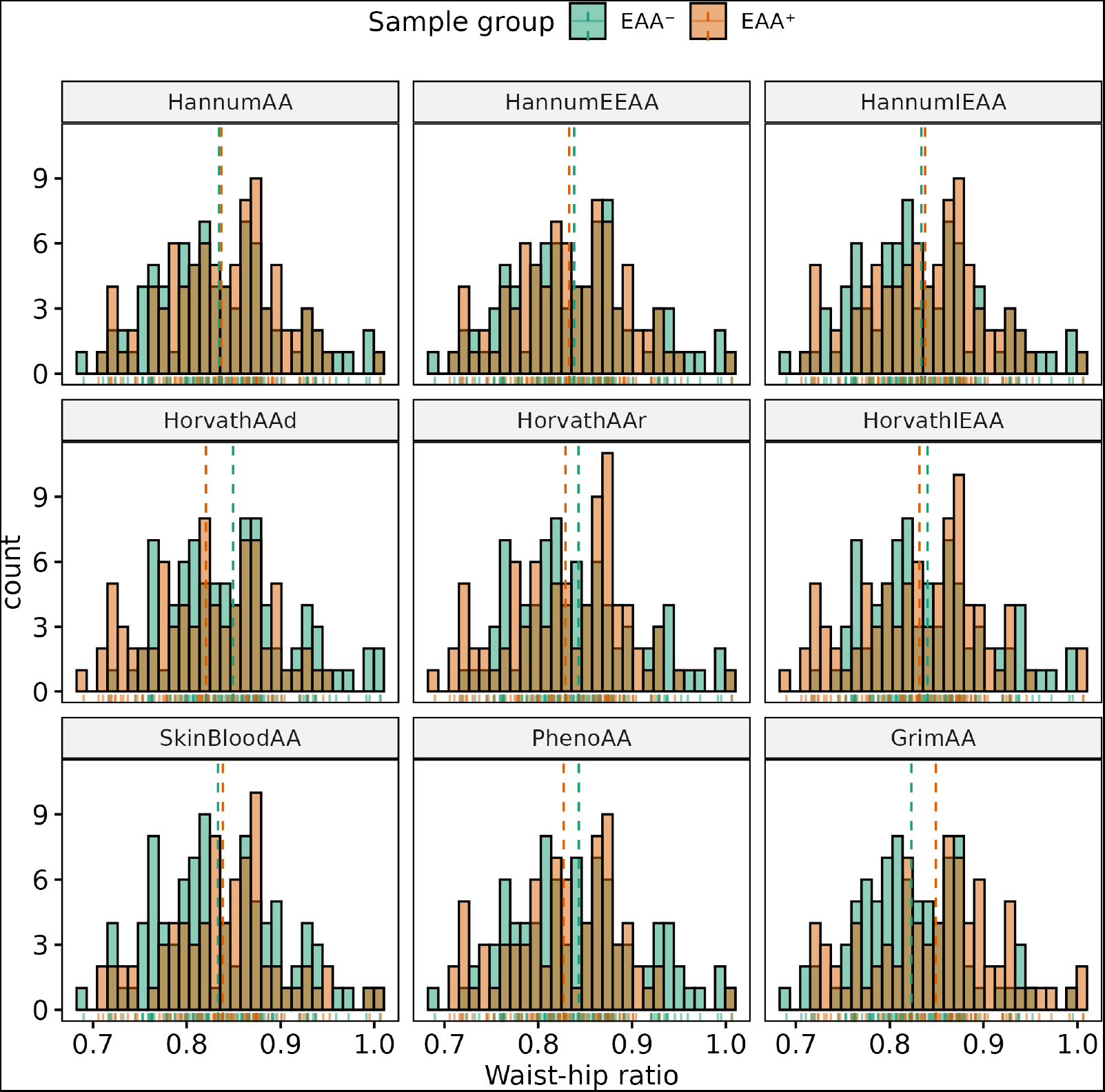
Histograms of distributions of waist-hip ratio in females in EAA^+^ and EAA^−^ groups. Dashed lines correspond to the group means.

**Figure A7.**
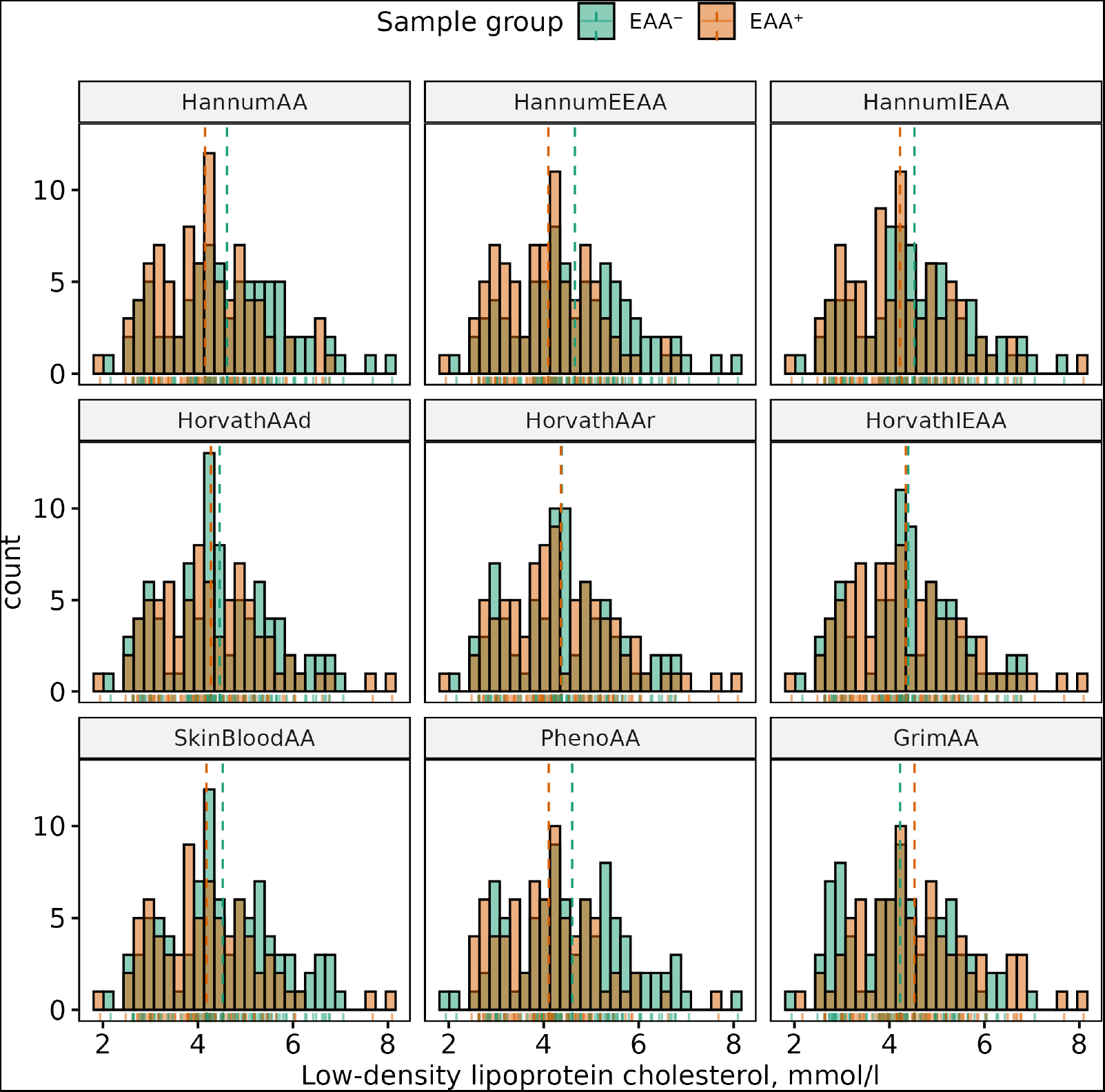
Histograms of distributions of LDL levels in females in EAA^+^ and EAA^−^ groups. Dashed lines correspond to the group means.

**Figure A8.**
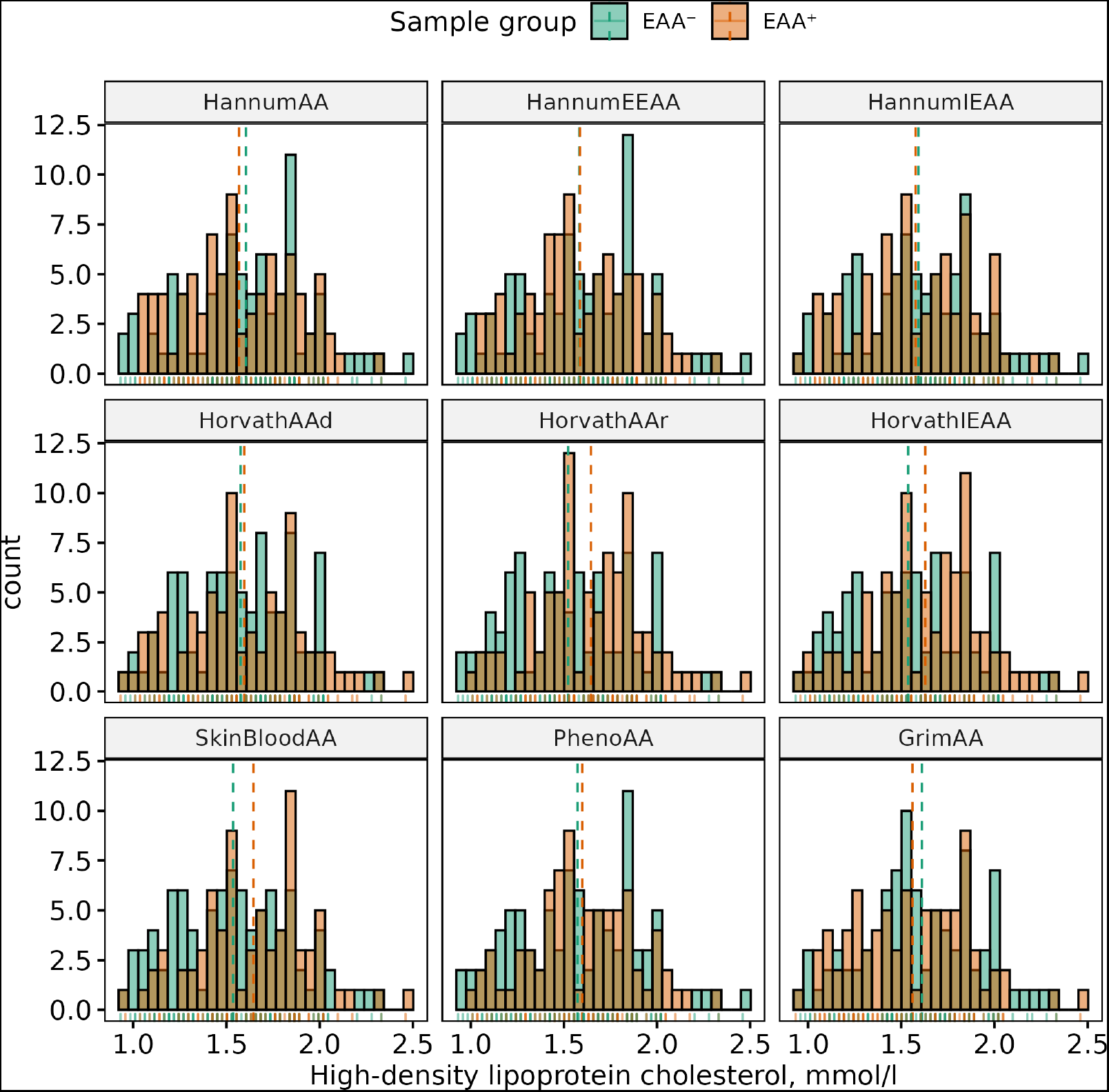
Histograms of distributions of HDL levels in females in EAA^+^ and EAA^−^ groups. Dashed lines correspond to the group means.

**Figure A9.**
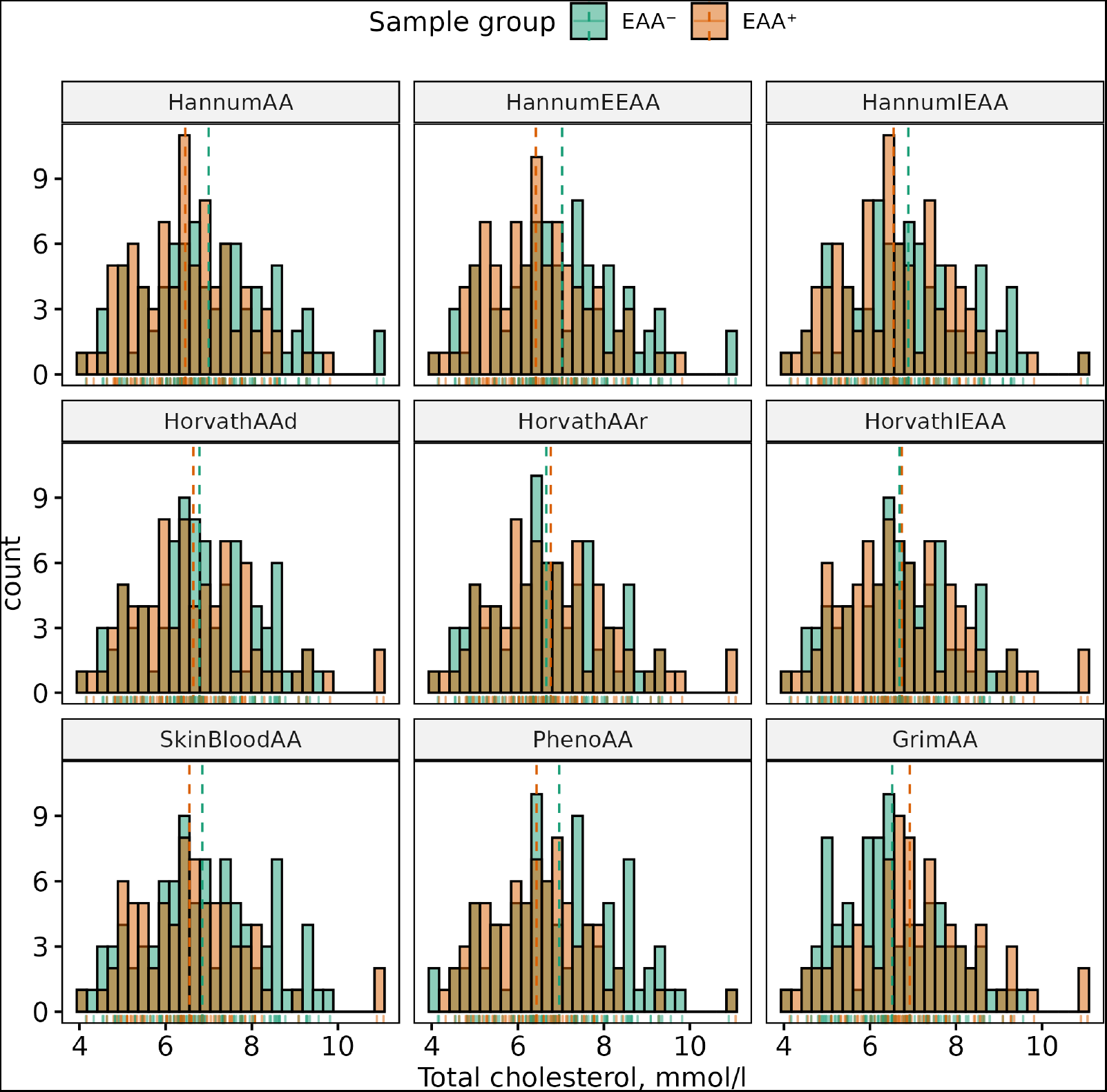
Histograms of distributions of TC levels in females in EAA^+^ and EAA^−^ groups. Dashed lines correspond to the group means.

**Figure A10.**
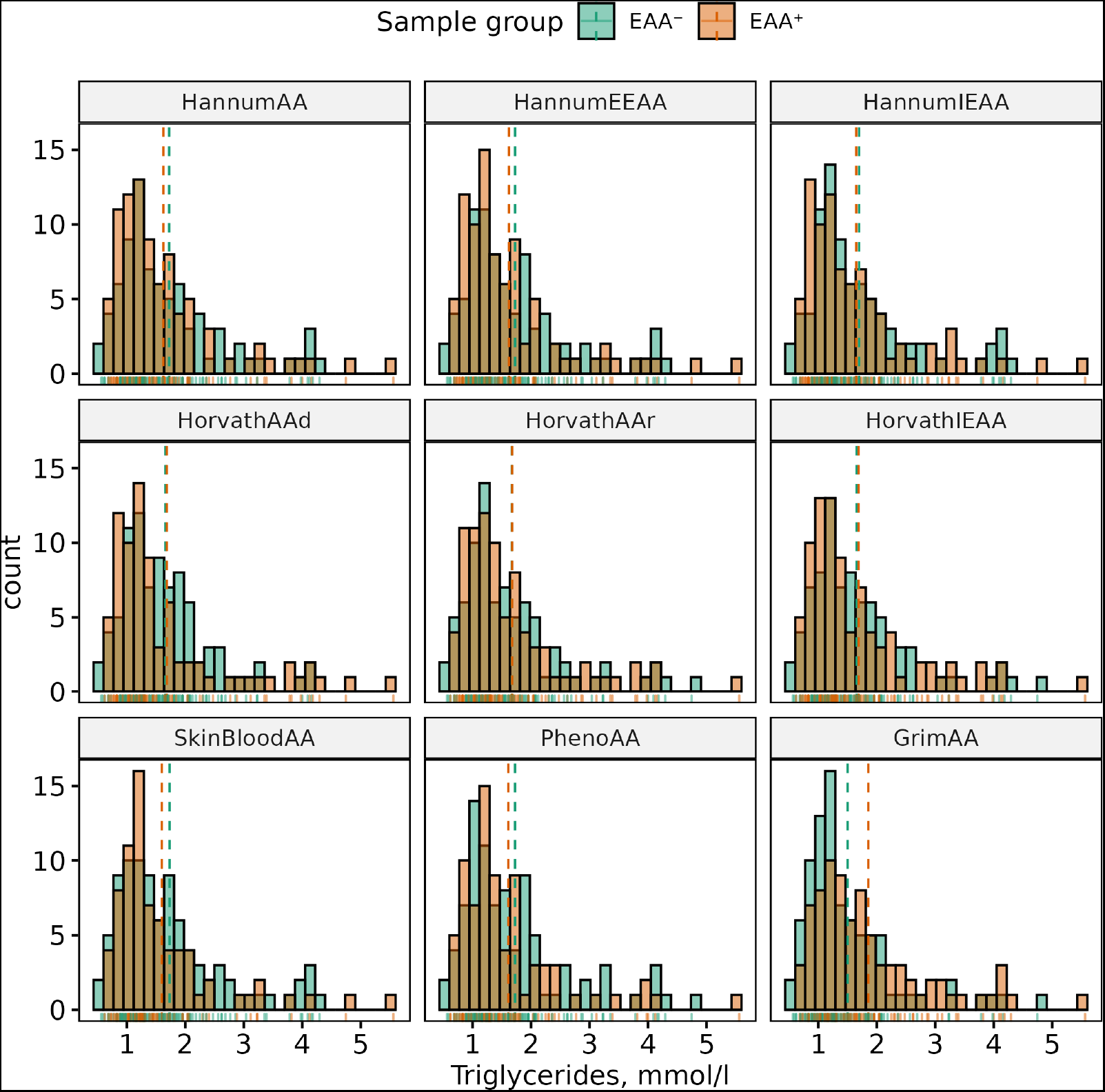
Histograms of distributions of TG levels in females in EAA^+^ and EAA^−^ groups. Dashed lines correspond to the group means.

**Figure A11.**
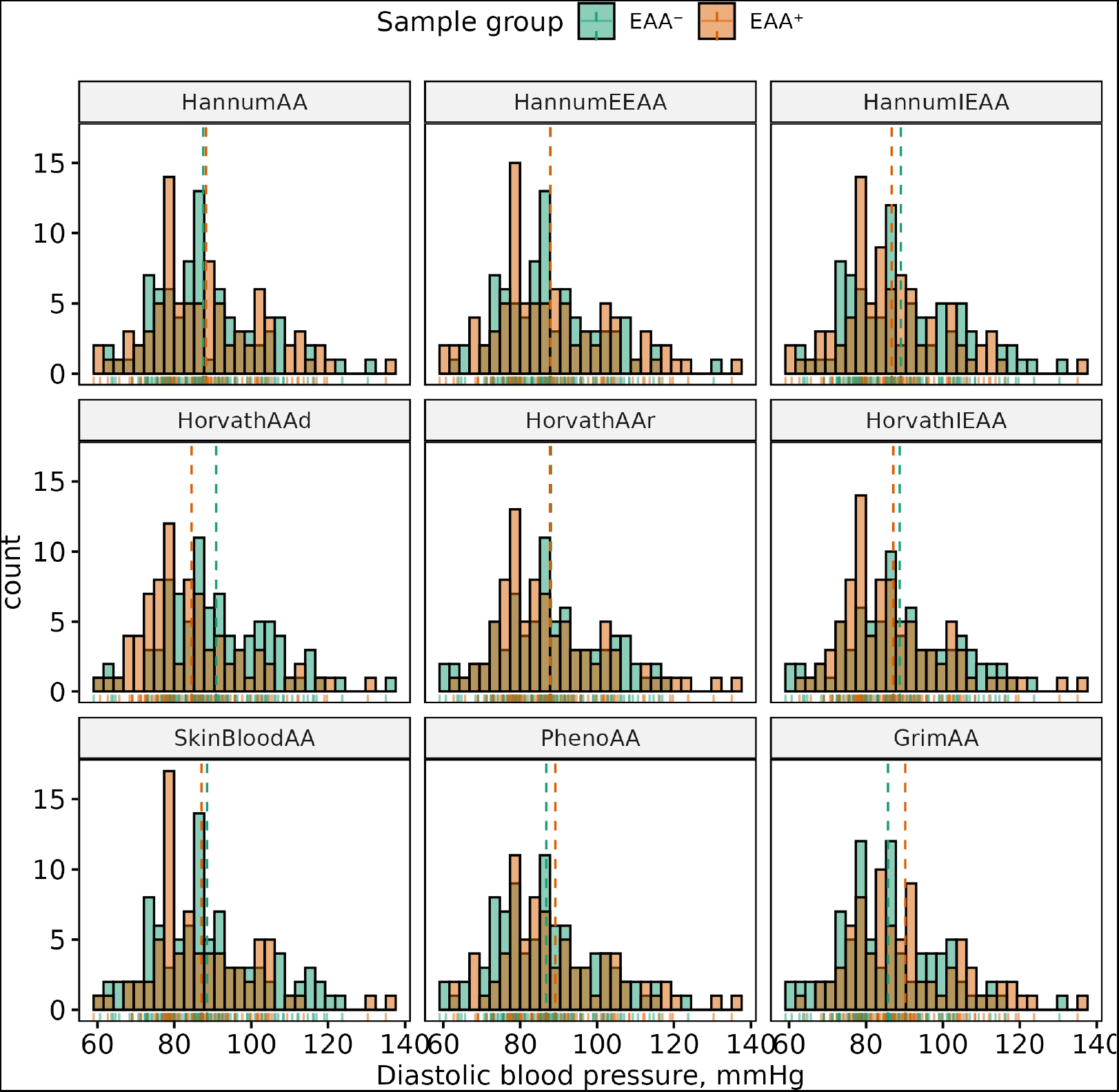
Histograms of distributions of DBP levels in females in EAA^+^ and EAA^−^ groups. Dashed lines correspond to the group means.

**Figure A12.**
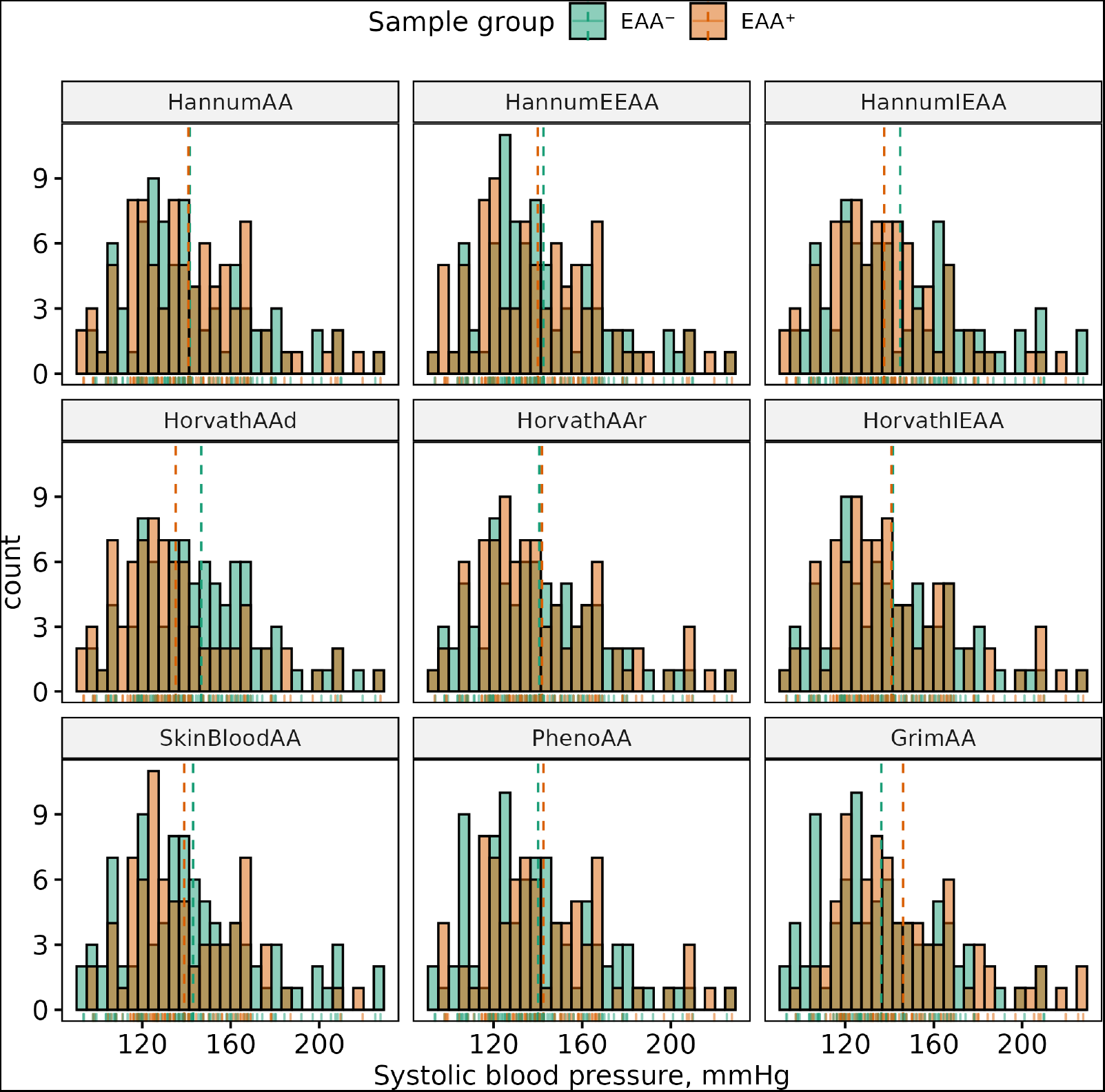
Histograms of distributions of SBP levels in females in EAA^+^ and EAA^−^ groups. Dashed lines correspond to the group means.

**Table A3.** Summary of binary phenotypes. Being case means having phenotype. Smoking status cases correspond to current or former smokers. *p*-values and odds ratios (with corresponding 95% confidence intervals) obtained from the Fisher’s exact test testing difference in ratio of each class of the variable between male and female groups (**H**_0_: Classification for given binary variable for male and female groups are not different).

| Phenotype | Sex | Samples, n | Cases, n (%) | OR (95%CI) | p-value |
| --- | --- | --- | --- | --- | --- |
| Smoking status | All | 306 | 126 (41.18%) |  | < 0.001 |
|  | F | 166 | 24 (14.46%) | 0.06 (0.03, 0.12) |  |
|  | M | 140 | 102 (72.86%) | 15.69 (8.67, 29.34) |  |
| T2DM | All | 306 | 34 (11.11%) |  | 0.466 |
|  | F | 166 | 16 (9.64%) | 0.72 (0.33, 1.57) |  |
|  | M | 140 | 18 (12.86%) | 1.38 (0.64, 3.03) |  |
| CHD | All | 306 | 130 (42.48%) |  | 0.083 |
|  | F | 166 | 63 (37.95%) | 0.67 (0.41, 1.08) |  |
|  | M | 140 | 67 (47.86%) | 1.5 (0.93, 2.43) |  |
| CP | All | 105 | 46 (43.81%) |  | 0.698 |
|  | F | 55 | 23 (41.82%) | 0.85 (0.36, 1.96) |  |
|  | M | 50 | 23 (46%) | 1.18 (0.51, 2.75) |  |
| MCP | All | 105 | 16 (15.24%) |  | 0.001 |
|  | F | 55 | 2 (3.64%) | 0.1 (0.01, 0.47) |  |
|  | M | 50 | 14 (28%) | 10.1 (2.12, 96.93) |  |
| HT | All | 306 | 176 (57.52%) |  | 0.643 |
|  | F | 166 | 93 (56.02%) | 0.88 (0.54, 1.41) |  |
|  | M | 140 | 83 (59.29%) | 1.14 (0.71, 1.85) |  |

**Table A4.**
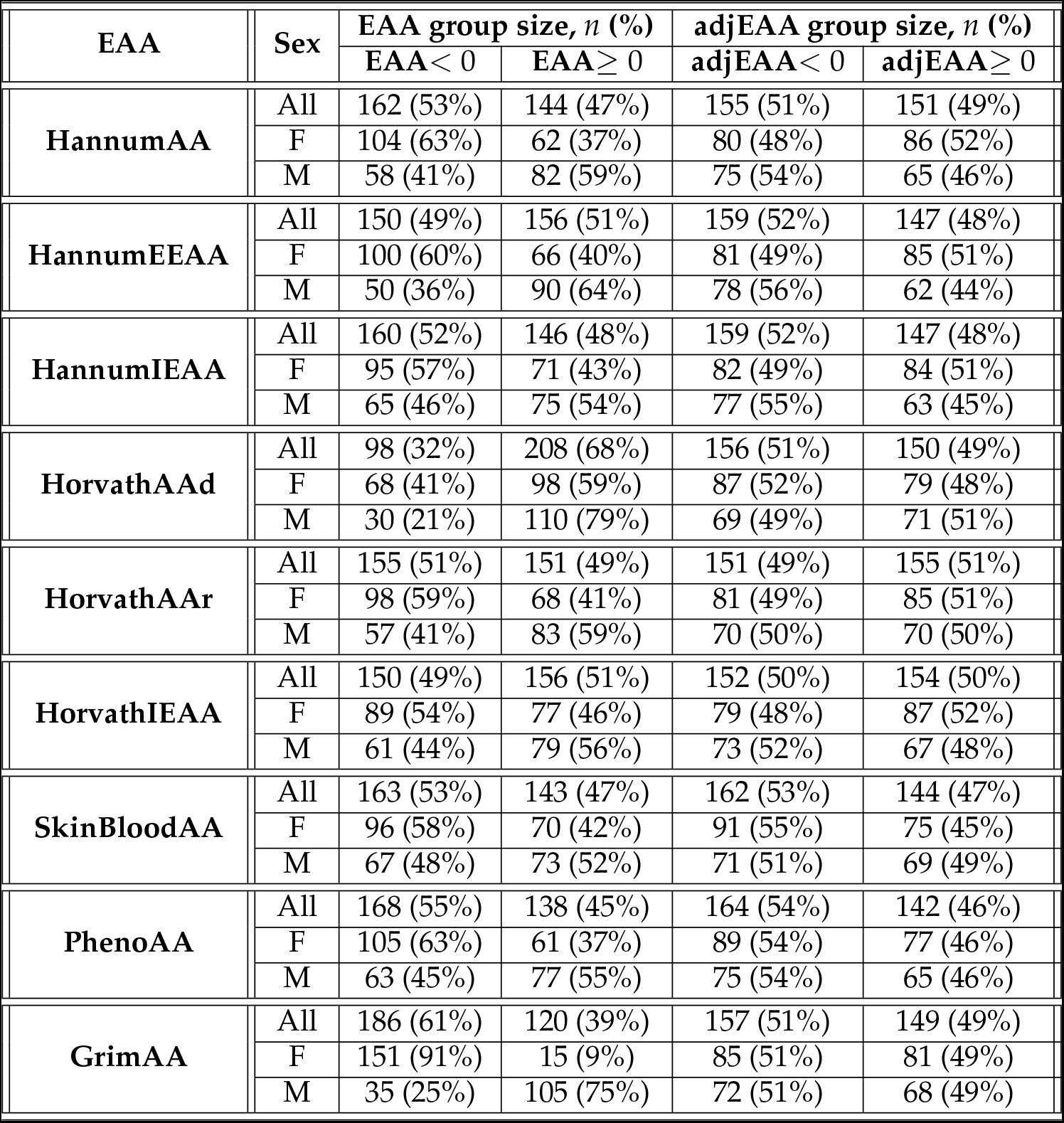
Number of samples in EAA^−^ and EAA^+^ groups for unadjusted and adjusted for sex EAA scores.

**Table A5.**
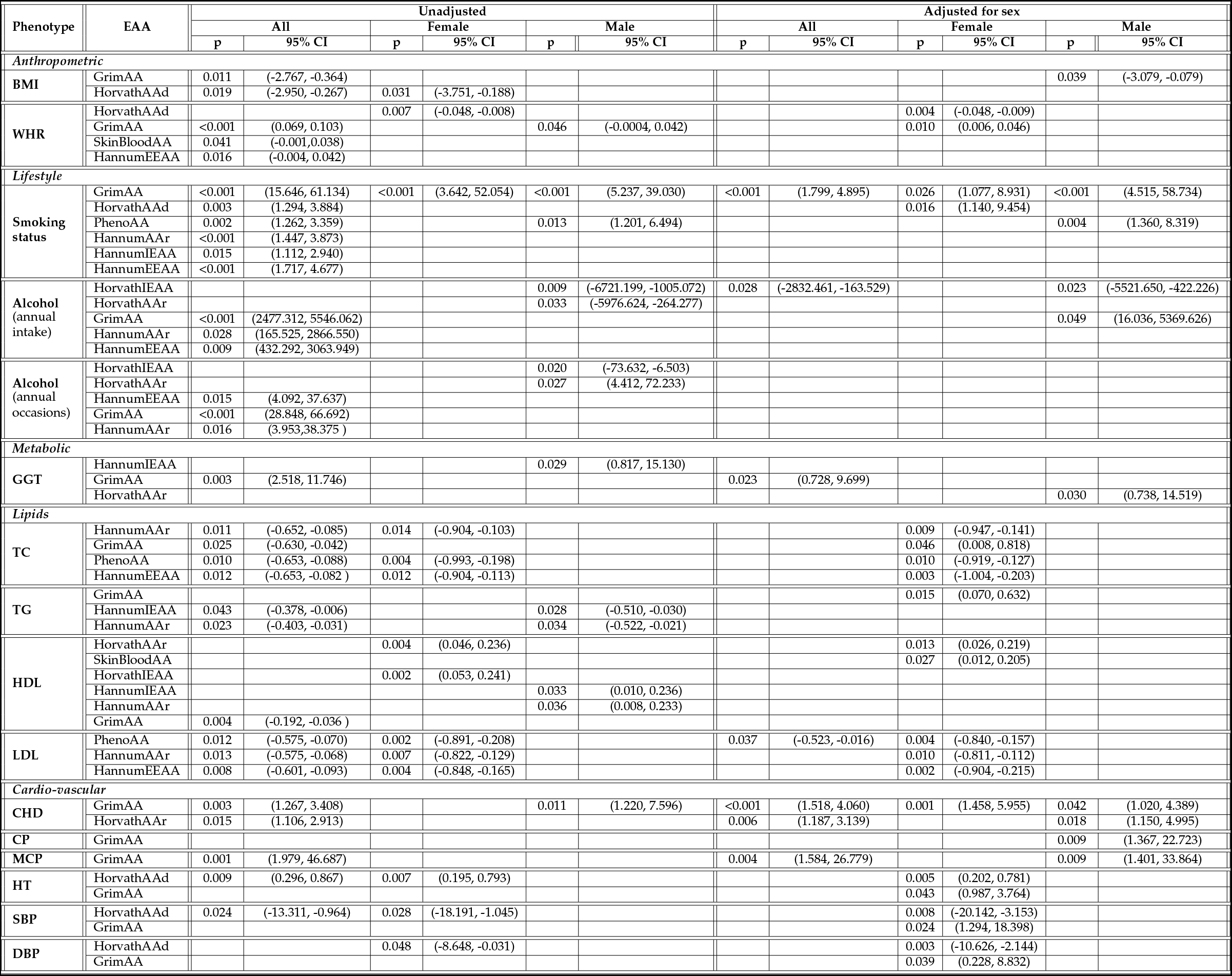
Significant differences in EAA^−^ and EAA^+^ groups for unadjusted and adjusted for sex EAA scores.

**Table A6.** All results for EAA^−^ and EAA^+^ groups for adjusted for sex EAA scores. For Welch’s *t*-test group 1 is *EAA*^−^, group 2 is *EAA*^+^. For Fisher’s exact test odds ratios depend on the phenotype value and grouping for the first sample in our data.

| Phenotype | EAA | p, all | Δmean/OR (95% CI, all) | p, female | Δmean/OR (95% CI, female) | p, male | Δmean/OR (95% CI, male) |
| --- | --- | --- | --- | --- | --- | --- | --- |
| ***Anthropometric*** |  |  |  |  |  |  |  |
| BMI | HorvathAAd | 0.118 | 0.946 (-0.242, 2.133) | 0.051 | 1.712 (-0.005, 3.428) | 0.880 | -0.117 (-1.641, 1.408) |
| BMI | HorvathAAr | 0.955 | 0.034 (-1.159, 1.227) | 0.665 | -0.381 (-2.116, 1.353) | 0.446 | 0.588 (-0.933, 2.108) |
| BMI | SkinBloodAA | 0.768 | -0.179 (-1.373, 1.014) | 0.873 | 0.141 (-1.593, 1.875) | 0.323 | -0.764 (-2.285, 0.758) |
| BMI | HorvathIEAA | 0.409 | 0.501 (-0.692, 1.694) | 0.742 | 0.291 (-1.452, 2.034) | 0.199 | 0.983 (-0.522, 2.488) |
| BMI | HannumEEAA | 0.691 | 0.242 (-0.958, 1.442) | 0.970 | -0.033 (-1.768, 1.702) | 0.228 | 0.929 (-0.589, 2.447) |
| BMI | HannumIEAA | 0.465 | 0.442 (-0.746, 1.631) | 0.656 | 0.394 (-1.348, 2.135) | 0.296 | 0.789 (-0.698, 2.277) |
| BMI | HannumAA | 0.982 | 0.014 (-1.181, 1.208) | 0.830 | 0.190 (-1.556, 1.935) | 0.921 | 0.078 (-1.460, 1.615) |
| BMI | GrimAA | 0.844 | -0.120 (-1.322, 1.082) | 0.081 | -1.543 (-3.281, 0.195) | 0.039 | 1.579 (0.079, 3.079) |
| BMI | PhenoAA | 0.841 | -0.123 (-1.329, 1.084) | 0.787 | 0.239 (-1.510, 1.989) | 0.483 | -0.554 (-2.113, 1.005) |
| WHR | HorvathAAd | 0.178 | 0.013 (-0.006, 0.032) | 0.004 | 0.029 (0.009, 0.048) | 0.909 | 0.001 (-0.021, 0.023) |
| WHR | HorvathAAr | 0.482 | 0.007 (-0.012, 0.026) | 0.170 | 0.014 (-0.006, 0.034) | 0.686 | -0.004 (-0.026, 0.017) |
| WHR | SkinBloodAA | 0.184 | -0.013 (-0.032, 0.006) | 0.604 | -0.005 (-0.025, 0.015) | 0.269 | -0.012 (-0.034, 0.010) |
| WHR | HorvathIEAA | 0.317 | 0.010 (-0.009, 0.028) | 0.376 | 0.009 (-0.011, 0.029) | 0.996 | 0.000 (-0.022, 0.022) |
| WHR | HannumEEAA | 0.152 | 0.014 (-0.005, 0.033) | 0.605 | 0.005 (-0.015, 0.025) | 0.477 | 0.008 (-0.014, 0.030) |
| WHR | HannumIEAA | 0.506 | 0.006 (-0.012, 0.025) | 0.678 | -0.004 (-0.024, 0.016) | 0.583 | 0.006 (-0.016, 0.028) |
| WHR | HannumAA | 0.609 | 0.005 (-0.014, 0.024) | 0.810 | -0.002 (-0.023, 0.018) | 0.911 | 0.001 (-0.021, 0.023) |
| WHR | GrimAA | 0.162 | -0.013 (-0.032, 0.005) | 0.010 | -0.026 (-0.046, -0.006) | 0.920 | 0.001 (-0.021, 0.023) |
| WHR | PhenoAA | 0.475 | 0.007 (-0.012, 0.026) | 0.104 | 0.016 (-0.003, 0.036) | 0.714 | -0.004 (-0.026, 0.018) |
| ***Lifestyle*** |  |  |  |  |  |  |  |
| Alcohol (annual intake) | HorvathAAd | 0.496 | 459.390 (-868.299, 1787.078) | 0.292 | -225.055 (-645.973, 195.864) | 0.238 | 1595.070 (-1070.824, 4260.965) |
| Alcohol (annual intake) | HorvathAAr | 0.107 | 1101.018 (-238.071, 2440.106) | 0.522 | -130.099 (-531.505, 271.307) | 0.069 | 2435.000 (-192.278, 5062.278) |
| Alcohol (annual intake) | SkinBloodAA | 0.438 | 519.830 (-797.085, 1836.746) | 0.388 | -190.837 (-627.538, 245.865) | 0.182 | 1783.619 (-845.651, 4412.889) |
| Alcohol (annual intake) | HorvathIEAA | 0.028 | 1497.995 (163.529, 2832.461) | 0.485 | -140.131 (-535.868, 255.606) | 0.023 | 2971.938 (422.226, 5521.650) |
| Alcohol (annual intake) | HannumEEAA | 0.833 | 144.007 (-1197.565, 1485.579) | 0.373 | -180.731 (-581.352, 219.891) | 0.888 | -193.486 (-2906.095, 2519.122) |
| Alcohol (annual intake) | HannumIEAA | 0.506 | 448.790 (-878.267, 1775.846) | 0.379 | -180.038 (-583.961, 223.885) | 0.642 | 616.522 (-2000.890, 3233.935) |
| Alcohol (annual intake) | HannumAA | 0.529 | 426.629 (-906.939, 1760.197) | 0.257 | -228.406 (-625.223, 168.412) | 0.627 | 646.862 (-1977.101, 3270.824) |
| Alcohol (annual intake) | GrimAA | 0.105 | -1119.352 (-2474.041, 235.337) | 0.355 | 187.723 (-212.549, 587.995) | 0.049 | -2692.831 (-5369.626, -16.036) |
| Alcohol (annual intake) | PhenoAA | 0.446 | 510.560 (-806.875, 1827.995) | 0.727 | -76.111 (-506.575, 354.353) | 0.360 | 1210.595 (-1395.209, 3816.398) |
| Alcohol (annual occasions) | HorvathAAd | 0.886 | -1.236 (-18.230, 15.759) | 0.316 | -5.429 (-16.092, 5.234) | 0.629 | 7.843 (-24.213, 39.900) |
| Alcohol (annual occasions) | HorvathAAr | 0.213 | 10.806 (-6.247, 27.859) | 0.441 | -4.114 (-14.640, 6.412) | 0.096 | 26.900 (-4.801, 58.601) |
| Alcohol (annual occasions) | SkinBloodAA | 0.884 | 1.253 (-15.668, 18.175) | 0.375 | -4.829 (-15.541, 5.883) | 0.392 | 13.818 (-18.028, 45.664) |
| Alcohol (annual occasions) | HorvathIEAA | 0.134 | 13.001 (-4.012, 30.013) | 0.419 | -4.305 (-14.803, 6.194) | 0.083 | 27.554 (-3.661, 58.768) |
| Alcohol (annual occasions) | HannumEEAA | 0.967 | -0.360 (-17.423, 16.703) | 0.515 | -3.487 (-14.028, 7.054) | 0.722 | -5.854 (-38.391, 26.683) |
| Alcohol (annual occasions) | HannumIEAA | 0.907 | -1.014 (-18.093, 16.065) | 0.587 | -2.911 (-13.470, 7.647) | 0.706 | -6.212 (-38.690, 26.266) |
| Alcohol (annual occasions) | HannumAA | 0.942 | 0.634 (-16.385, 17.654) | 0.255 | -6.066 (-16.547, 4.414) | 0.927 | 1.484 (-30.599, 33.568) |
| Alcohol (annual occasions) | GrimAA | 0.737 | -2.923 (-20.017, 14.170) | 0.529 | 3.366 (-7.167, 13.900) | 0.512 | -10.678 (-42.810, 21.453) |
| Alcohol (annual occasions) | PhenoAA | 0.412 | 7.037 (-9.830, 23.903) | 0.576 | 3.000 (-7.580, 13.580) | 0.459 | 11.880 (-19.746, 43.506) |
| Smoking status | HorvathAAd | 0.164 | 1.400 (0.865, 2.271) | 0.016 | 3.112 (1.140, 9.454) | 0.850 | 0.901 (0.398, 2.028) |
| Smoking status | HorvathAAr | 0.908 | 0.957 (0.591, 1.547) | 0.661 | 0.779 (0.294, 2.027) | 0.849 | 1.154 (0.513, 2.613) |
| Smoking status | SkinBloodAA | 0.908 | 1.039 (0.642, 1.682) | 0.826 | 0.847 (0.314, 2.209) | 1.000 | 0.962 (0.426, 2.169) |
| Smoking status | HorvathIEAA | 1.000 | 0.978 (0.605, 1.582) | 1.000 | 1.085 (0.417, 2.874) | 0.706 | 1.186 (0.527, 2.700) |
| Smoking status | HannumEEAA | 0.642 | 1.142 (0.706, 1.849) | 0.512 | 1.397 (0.536, 3.769) | 0.181 | 1.771 (0.771, 4.217) |
| Smoking status | HannumIEAA | 0.642 | 1.142 (0.706, 1.849) | 0.271 | 1.757 (0.669, 4.873) | 0.451 | 1.360 (0.600, 3.150) |
| Smoking status | HannumAA | 0.296 | 1.297 (0.802, 2.102) | 0.128 | 2.048 (0.769, 5.900) | 0.186 | 1.708 (0.750, 4.004) |
| Smoking status | GrimAA | 0.000 | 2.954 (1.799, 4.895) | 0.026 | 2.941 (1.077, 8.931) | 0.000 | 14.047 (4.515, 58.734) |
| Smoking status | PhenoAA | 0.132 | 1.424 (0.879, 2.311) | 0.663 | 0.801 (0.297, 2.087) | 0.004 | 3.249 (1.360, 8.319) |
| **Metabolic** |  |  |  |  |  |  |  |
| T2DM | HorvathAAd | 0.717 | 1.192 (0.548, 2.614) | 0.441 | 0.635 (0.180, 2.043) | 0.207 | 2.124 (0.685, 7.363) |
| T2DM | HorvathAAr | 0.102 | 1.912 (0.866, 4.419) | 0.794 | 1.250 (0.391, 4.172) | 0.075 | 2.943 (0.914, 11.207) |
| T2DM | SkinBloodAA | 0.720 | 1.141 (0.523, 2.489) | 0.188 | 2.169 (0.673, 7.658) | 0.451 | 0.618 (0.190, 1.882) |
| T2DM | HorvathIEAA | 0.364 | 1.469 (0.674, 3.284) | 0.798 | 1.186 (0.371, 3.956) | 0.313 | 1.844 (0.605, 6.012) |
| T2DM | HannumEEAA | 0.468 | 0.732 (0.328, 1.596) | 0.794 | 1.250 (0.391, 4.172) | 0.203 | 0.441 (0.116, 1.420) |
| T2DM | HannumIEAA | 1.000 | 0.957 (0.436, 2.081) | 0.432 | 1.706 (0.530, 6.018) | 0.321 | 0.572 (0.165, 1.777) |
| T2DM | HannumAA | 1.000 | 1.030 (0.472, 2.245) | 0.436 | 1.618 (0.503, 5.706) | 0.615 | 0.704 (0.216, 2.144) |
| T2DM | GrimAA | 0.858 | 0.929 (0.424, 2.021) | 1.000 | 1.054 (0.326, 3.410) | 0.803 | 0.828 (0.264, 2.510) |
| T2DM | PhenoAA | 0.586 | 0.788 (0.353, 1.718) | 0.112 | 0.354 (0.079, 1.235) | 0.455 | 1.518 (0.501, 4.757) |
| GGT | HorvathAAd | 0.664 | 0.990 (-3.484, 5.464) | 0.260 | 3.176 (-2.373, 8.725) | 0.770 | -1.037 (-8.046, 5.973) |
| GGT | HorvathAAr | 0.614 | -1.150 (-5.630, 3.330) | 0.151 | 4.135 (-1.521, 9.791) | 0.030 | -7.629 (-14.519, -0.738) |
| GGT | SkinBloodAA | 0.531 | 1.431 (-3.061, 5.924) | 0.794 | -0.766 (-6.551, 5.019) | 0.178 | 4.752 (-2.182, 11.685) |
| GGT | HorvathIEAA | 0.937 | 0.180 (-4.309, 4.669) | 0.305 | 2.983 (-2.749, 8.715) | 0.270 | -3.955 (-11.026, 3.115) |
| GGT | HannumEEAA | 0.186 | 2.997 (-1.449, 7.443) | 0.192 | 3.749 (-1.902, 9.400) | 0.803 | 0.873 (-6.041, 7.788) |
| GGT | HannumIEAA | 0.218 | 2.774 (-1.652, 7.201) | 0.893 | 0.386 (-5.266, 6.037) | 0.172 | 4.644 (-2.039, 11.326) |
| GGT | HannumAA | 0.120 | 3.530 (-0.925, 7.984) | 0.304 | 2.968 (-2.716, 8.651) | 0.344 | 3.255 (-3.530, 10.041) |
| GGT | GrimAA | 0.023 | -5.213 (-9.699, -0.728) | 0.087 | -4.906 (-10.542, 0.729) | 0.114 | -5.618 (-12.606, 1.370) |
| GGT | PhenoAA | 0.916 | 0.244 (-4.285, 4.773) | 0.205 | 3.549 (-1.960, 9.057) | 0.314 | -3.666 (-10.841, 3.510) |
| Glucose | HorvathAAd | 0.500 | 0.130 (-0.248, 0.508) | 0.197 | 0.325 (-0.170, 0.820) | 0.761 | -0.089 (-0.664, 0.487) |
| Glucose | HorvathAAr | 0.706 | -0.072 (-0.447, 0.303) | 0.907 | -0.029 (-0.527, 0.468) | 0.675 | -0.123 (-0.701, 0.455) |
| Glucose | SkinBloodAA | 0.686 | 0.077 (-0.297, 0.452) | 0.778 | -0.072 (-0.572, 0.428) | 0.392 | 0.249 (-0.325, 0.824) |
| Glucose | HorvathIEAA | 0.873 | -0.031 (-0.407, 0.346) | 0.981 | -0.006 (-0.503, 0.491) | 0.828 | -0.065 (-0.653, 0.524) |
| Glucose | HannumEEAA | 0.455 | 0.143 (-0.234, 0.521) | 0.910 | -0.029 (-0.533, 0.476) | 0.260 | 0.335 (-0.251, 0.921) |
| Glucose | HannumIEAA | 0.360 | -0.182 (-0.574, 0.209) | 0.088 | -0.439 (-0.944, 0.067) | 0.730 | 0.108 (-0.509, 0.725) |
| Glucose | HannumAA | 0.488 | -0.133 (-0.512, 0.245) | 0.564 | -0.147 (-0.647, 0.354) | 0.680 | -0.128 (-0.742, 0.486) |
| Glucose | GrimAA | 0.792 | 0.051 (-0.328, 0.430) | 0.359 | -0.234 (-0.736, 0.268) | 0.200 | 0.376 (-0.201, 0.952) |
| Glucose | PhenoAA | 0.517 | 0.124 (-0.253, 0.501) | 0.421 | 0.203 (-0.294, 0.700) | 0.909 | 0.034 (-0.550, 0.618) |
| **Lipids** |  |  |  |  |  |  |  |
| HDL | HorvathAAd | 0.948 | 0.002 (-0.073, 0.077) | 0.712 | -0.018 (-0.117, 0.080) | 0.725 | 0.021 (-0.096, 0.137) |
| HDL | HorvathAAr | 0.281 | -0.041 (-0.116, 0.034) | 0.013 | -0.122 (-0.219, -0.026) | 0.328 | 0.057 (-0.058, 0.173) |
| HDL | SkinBloodAA | 0.104 | -0.062 (-0.136, 0.013) | 0.027 | -0.108 (-0.205, -0.012) | 0.790 | -0.016 (-0.131, 0.100) |
| HDL | HorvathIEAA | 0.328 | -0.037 (-0.113, 0.038) | 0.065 | -0.091 (-0.188, 0.006) | 0.541 | 0.035 (-0.079, 0.150) |
| HDL | HannumEEAA | 0.158 | -0.054 (-0.129, 0.021) | 0.959 | -0.003 (-0.101, 0.095) | 0.092 | -0.102 (-0.221, 0.017) |
| HDL | HannumIEAA | 0.195 | -0.050 (-0.125, 0.026) | 0.765 | 0.015 (-0.083, 0.113) | 0.056 | -0.116 (-0.234, 0.003) |
| HDL | HannumAA | 0.529 | -0.024 (-0.099, 0.051) | 0.450 | 0.038 (-0.060, 0.135) | 0.148 | -0.087 (-0.204, 0.031) |
| HDL | GrimAA | 0.935 | -0.003 (-0.078, 0.072) | 0.306 | 0.051 (-0.047, 0.148) | 0.260 | -0.066 (-0.183, 0.050) |
| HDL | PhenoAA | 0.604 | -0.020 (-0.095, 0.055) | 0.602 | -0.026 (-0.122, 0.071) | 0.825 | -0.013 (-0.129, 0.103) |
| LDL | HorvathAAd | 0.458 | 0.097 (-0.160, 0.353) | 0.324 | 0.177 (-0.176, 0.531) | 0.916 | -0.020 (-0.388, 0.349) |
| LDL | HorvathAAr | 0.643 | -0.060 (-0.316, 0.195) | 0.916 | 0.019 (-0.334, 0.372) | 0.435 | -0.146 (-0.515, 0.223) |
| LDL | SkinBloodAA | 0.138 | 0.193 (-0.063, 0.449) | 0.055 | 0.343 (-0.007, 0.693) | 0.954 | -0.011 (-0.381, 0.359) |
| LDL | HorvathIEAA | 0.627 | -0.063 (-0.319, 0.193) | 0.779 | 0.050 (-0.302, 0.402) | 0.379 | -0.166 (-0.539, 0.207) |
| LDL | HannumEEAA | 0.096 | 0.216 (-0.039, 0.470) | 0.002 | 0.560 (0.215, 0.904) | 0.441 | -0.146 (-0.520, 0.228) |
| LDL | HannumIEAA | 0.427 | 0.104 (-0.153, 0.360) | 0.090 | 0.303 (-0.048, 0.654) | 0.617 | -0.095 (-0.468, 0.279) |
| LDL | HannumAA | 0.211 | 0.163 (-0.093, 0.418) | 0.010 | 0.461 (0.112, 0.811) | 0.413 | -0.154 (-0.525, 0.217) |
| LDL | GrimAA | 0.446 | -0.099 (-0.356, 0.157) | 0.089 | -0.304 (-0.655, 0.047) | 0.439 | 0.145 (-0.224, 0.513) |
| LDL | PhenoAA | 0.037 | 0.270 (0.016, 0.523) | 0.004 | 0.498 (0.157, 0.840) | 0.994 | -0.001 (-0.373, 0.371) |
| TC | HorvathAAd | 0.560 | 0.085 (-0.203, 0.374) | 0.479 | 0.148 (-0.263, 0.558) | 0.920 | -0.020 (-0.411, 0.371) |
| TC | HorvathAAr | 0.410 | -0.120 (-0.407, 0.167) | 0.610 | -0.105 (-0.513, 0.302) | 0.525 | -0.126 (-0.517, 0.265) |
| TC | SkinBloodAA | 0.203 | 0.186 (-0.101, 0.472) | 0.149 | 0.297 (-0.108, 0.702) | 0.947 | 0.013 (-0.378, 0.404) |
| TC | HorvathIEAA | 0.404 | -0.122 (-0.409, 0.165) | 0.797 | -0.053 (-0.460, 0.353) | 0.431 | -0.157 (-0.552, 0.237) |
| TC | HannumEEAA | 0.140 | 0.215 (-0.071, 0.500) | 0.003 | 0.603 (0.203, 1.004) | 0.378 | -0.178 (-0.575, 0.220) |
| TC | HannumIEAA | 0.473 | 0.105 (-0.182, 0.392) | 0.099 | 0.341 (-0.064, 0.747) | 0.552 | -0.120 (-0.516, 0.277) |
| TC | HannumAA | 0.192 | 0.190 (-0.096, 0.477) | 0.009 | 0.544 (0.141, 0.947) | 0.385 | -0.174 (-0.568, 0.220) |
| TC | GrimAA | 0.215 | -0.182 (-0.470, 0.106) | 0.046 | -0.413 (-0.818, -0.008) | 0.633 | 0.095 (-0.296, 0.485) |
| TC | PhenoAA | 0.081 | 0.253 (-0.031, 0.538) | 0.010 | 0.523 (0.127, 0.919) | 0.738 | -0.067 (-0.460, 0.326) |
| TG | HorvathAAd | 0.751 | -0.031 (-0.220, 0.159) | 0.867 | -0.025 (-0.314, 0.265) | 0.702 | -0.046 (-0.285, 0.192) |
| TG | HorvathAAr | 0.667 | -0.041 (-0.229, 0.147) | 0.979 | -0.004 (-0.288, 0.280) | 0.498 | -0.082 (-0.322, 0.157) |
| TG | SkinBloodAA | 0.215 | 0.119 (-0.069, 0.308) | 0.342 | 0.137 (-0.148, 0.422) | 0.474 | 0.087 (-0.153, 0.327) |
| TG | HorvathIEAA | 0.623 | -0.047 (-0.235, 0.141) | 0.851 | -0.027 (-0.310, 0.256) | 0.633 | -0.059 (-0.302, 0.184) |
| TG | HannumEEAA | 0.222 | 0.117 (-0.071, 0.305) | 0.479 | 0.102 (-0.182, 0.386) | 0.198 | 0.155 (-0.082, 0.393) |
| TG | HannumIEAA | 0.243 | 0.112 (-0.076, 0.300) | 0.718 | 0.052 (-0.232, 0.336) | 0.097 | 0.200 (-0.036, 0.435) |
| TG | HannumAA | 0.233 | 0.114 (-0.074, 0.302) | 0.491 | 0.099 (-0.185, 0.383) | 0.222 | 0.147 (-0.090, 0.385) |
| TG | GrimAA | 0.070 | -0.174 (-0.363, 0.014) | 0.015 | -0.351 (-0.632, -0.070) | 0.767 | 0.036 (-0.204, 0.276) |
| TG | PhenoAA | 0.940 | 0.007 (-0.183, 0.198) | 0.445 | 0.110 (-0.174, 0.395) | 0.357 | -0.115 (-0.361, 0.131) |
| *Cardio-vascular* |  |  |  |  |  |  |  |
| CHD | HorvathAAd | 0.419 | 0.820 (0.507, 1.323) | 0.423 | 0.737 (0.372, 1.448) | 0.866 | 0.895 (0.437, 1.830) |
| CHD | HorvathAAr | 0.006 | 1.924 (1.187, 3.139) | 0.151 | 1.626 (0.826, 3.234) | 0.018 | 2.377 (1.150, 4.995) |
| CHD | SkinBloodAA | 1.000 | 0.991 (0.613, 1.600) | 0.521 | 0.776 (0.390, 1.528) | 0.612 | 1.253 (0.613, 2.572) |
| CHD | HorvathIEAA | 0.134 | 1.421 (0.880, 2.303) | 0.423 | 1.357 (0.690, 2.687) | 0.236 | 1.568 (0.765, 3.239) |
| CHD | HannumEEAA | 0.420 | 1.209 (0.749, 1.954) | 0.264 | 1.466 (0.746, 2.905) | 1.000 | 1.039 (0.505, 2.136) |
| CHD | HannumIEAA | 0.730 | 1.086 (0.673, 1.754) | 1.000 | 1.012 (0.516, 1.989) | 0.611 | 1.237 (0.603, 2.547) |
| CHD | HannumAA | 0.298 | 1.296 (0.803, 2.095) | 0.201 | 1.564 (0.795, 3.109) | 0.865 | 1.107 (0.540, 2.273) |
| CHD | GrimAA | 0.000 | 2.474 (1.518, 4.060) | 0.001 | 2.915 (1.458, 5.955) | 0.042 | 2.102 (1.020, 4.389) |
| CHD | PhenoAA | 0.418 | 1.218 (0.754, 1.969) | 0.873 | 1.083 (0.550, 2.128) | 0.397 | 1.393 (0.680, 2.870) |
| MCP | HorvathAAd | 0.429 | 0.638 (0.184, 2.119) | 1.000 | 0.964 (0.012, 78.414) | 0.343 | 0.485 (0.112, 1.986) |
| MCP | HorvathAAr | 0.290 | 1.797 (0.540, 6.237) | 0.170 | Inf (0.265, Inf) | 0.761 | 1.244 (0.302, 5.156) |
| MCP | SkinBloodAA | 0.426 | 0.616 (0.169, 2.062) | 1.000 | 1.298 (0.016, 105.663) | 0.211 | 0.404 (0.087, 1.672) |
| MCP | HorvathIEAA | 1.000 | 1.118 (0.333, 3.755) | 0.236 | Inf (0.196, Inf) | 1.000 | 0.841 (0.197, 3.436) |
| MCP | HannumEEAA | 1.000 | 1.092 (0.315, 3.640) | 1.000 | 1.400 (0.017, 114.021) | 1.000 | 1.049 (0.244, 4.318) |
| MCP | HannumIEAA | 0.412 | 0.535 (0.134, 1.842) | 1.000 | 1.118 (0.014, 90.951) | 0.353 | 0.507 (0.097, 2.191) |
| MCP | HannumAA | 1.000 | 1.043 (0.301, 3.475) | 1.000 | 1.298 (0.016, 105.663) | 1.000 | 1.049 (0.244, 4.318) |
| MCP | GrimAA | 0.004 | 5.795 (1.584, 26.779) | 0.156 | Inf (0.286, Inf) | 0.009 | 6.225 (1.401, 33.864) |
| MCP | PhenoAA | 0.791 | 1.144 (0.330, 3.815) | 1.000 | 1.400 (0.017, 114.021) | 1.000 | 1.175 (0.273, 4.866) |
| CP | HorvathAAd | 0.332 | 0.677 (0.289, 1.571) | 1.000 | 1.089 (0.328, 3.644) | 0.153 | 0.392 (0.104, 1.397) |
| CP | HorvathAAr | 0.843 | 1.141 (0.488, 2.666) | 1.000 | 1.122 (0.330, 3.793) | 1.000 | 1.143 (0.326, 4.030) |
| CP | SkinBloodAA | 0.555 | 0.746 (0.318, 1.730) | 1.000 | 0.989 (0.292, 3.317) | 0.395 | 0.536 (0.148, 1.873) |
| CP | HorvathIEAA | 0.437 | 1.380 (0.595, 3.225) | 0.176 | 2.239 (0.669, 7.862) | 0.782 | 0.832 (0.234, 2.910) |
| CP | HannumEEAA | 1.000 | 0.957 (0.406, 2.244) | 0.417 | 0.610 (0.171, 2.069) | 0.567 | 1.544 (0.435, 5.598) |
| CP | HannumIEAA | 1.000 | 0.977 (0.417, 2.278) | 1.000 | 1.038 (0.311, 3.462) | 1.000 | 0.936 (0.257, 3.359) |
| CP | HannumAA | 0.844 | 0.894 (0.380, 2.090) | 0.286 | 0.540 (0.151, 1.823) | 0.567 | 1.544 (0.435, 5.598) |
| CP | GrimAA | 0.074 | 2.090 (0.883, 5.032) | 1.000 | 0.941 (0.271, 3.199) | 0.009 | 5.241 (1.367, 22.723) |
| CP | PhenoAA | 1.000 | 1.026 (0.435, 2.412) | 0.787 | 0.829 (0.240, 2.796) | 0.774 | 1.301 (0.362, 4.717) |
| HT | HorvathAAd | 0.064 | 0.642 (0.396, 1.038) | 0.005 | 0.400 (0.202, 0.781) | 0.864 | 1.113 (0.537, 2.309) |
| HT | HorvathAAr | 0.908 | 1.046 (0.648, 1.689) | 0.877 | 0.941 (0.487, 1.818) | 0.731 | 1.193 (0.576, 2.480) |
| HT | SkinBloodAA | 0.563 | 0.860 (0.532, 1.389) | 0.351 | 0.744 (0.383, 1.440) | 1.000 | 1.011 (0.488, 2.097) |
| HT | HorvathIEAA | 0.908 | 0.970 (0.601, 1.565) | 0.876 | 0.930 (0.481, 1.797) | 1.000 | 1.033 (0.498, 2.146) |
| HT | HannumEEAA | 0.644 | 1.140 (0.706, 1.843) | 0.755 | 1.144 (0.592, 2.212) | 0.730 | 1.160 (0.557, 2.430) |
| HT | HannumIEAA | 0.817 | 1.081 (0.669, 1.746) | 0.876 | 1.096 (0.567, 2.118) | 0.864 | 1.080 (0.520, 2.256) |
| HT | HannumAA | 0.356 | 1.248 (0.773, 2.018) | 0.435 | 1.316 (0.681, 2.552) | 0.730 | 1.189 (0.573, 2.482) |
| HT | GrimAA | 0.488 | 1.193 (0.739, 1.929) | 0.043 | 1.917 (0.987, 3.764) | 0.303 | 0.677 (0.324, 1.403) |
| HT | PhenoAA | 0.908 | 0.965 (0.597, 1.559) | 0.876 | 1.088 (0.563, 2.109) | 0.610 | 0.834 (0.401, 1.731) |
| SBP | HorvathAAd | 0.069 | 5.357 (-0.419, 11.133) | 0.008 | 11.648 (3.153, 20.142) | 0.582 | -2.110 (-9.680, 5.461) |
| SBP | HorvathAAr | 0.606 | -1.520 (-7.316, 4.276) | 0.802 | -1.100 (-9.759, 7.559) | 0.602 | -2.005 (-9.585, 5.576) |
| SBP | SkinBloodAA | 0.506 | 1.949 (-3.810, 7.708) | 0.366 | 3.904 (-4.595, 12.403) | 0.920 | -0.386 (-7.976, 7.204) |
| SBP | HorvathIEAA | 0.931 | -0.256 (-6.057, 5.546) | 0.895 | 0.581 (-8.097, 9.260) | 0.756 | -1.198 (-8.816, 6.419) |
| SBP | HannumEEAA | 0.986 | -0.051 (-5.865, 5.763) | 0.570 | 2.492 (-6.159, 11.144) | 0.432 | -3.027 (-10.622, 4.569) |
| SBP | HannumIEAA | 0.356 | 2.712 (-3.058, 8.481) | 0.102 | 7.186 (-1.433, 15.805) | 0.505 | -2.565 (-10.151, 5.021) |
| SBP | HannumAA | 0.710 | -1.099 (-6.902, 4.704) | 0.884 | 0.641 (-8.019, 9.300) | 0.416 | -3.117 (-10.675, 4.441) |
| SBP | GrimAA | 0.293 | -3.104 (-8.899, 2.691) | 0.024 | -9.846 (-18.398, -1.294) | 0.200 | 4.894 (-2.621, 12.409) |
| SBP | PhenoAA | 0.793 | -0.779 (-6.619, 5.061) | 0.590 | -2.376 (-11.075, 6.322) | 0.774 | 1.114 (-6.549, 8.778) |
| DBP | HorvathAAd | 0.135 | 2.356 (-0.735, 5.448) | 0.003 | 6.385 (2.144, 10.626) | 0.309 | -2.288 (-6.718, 2.141) |
| DBP | HorvathAAr | 0.994 | -0.011 (-3.106, 3.084) | 0.922 | 0.215 (-4.131, 4.560) | 0.886 | -0.324 (-4.774, 4.127) |
| DBP | SkinBloodAA | 0.523 | 1.009 (-2.094, 4.113) | 0.520 | 1.416 (-2.924, 5.756) | 0.760 | 0.688 (-3.766, 5.142) |
| DBP | HorvathIEAA | 0.443 | 1.209 (-1.887, 4.305) | 0.462 | 1.626 (-2.730, 5.981) | 0.809 | 0.551 (-3.945, 5.047) |
| DBP | HannumEEAA | 0.870 | -0.259 (-3.374, 2.857) | 0.989 | 0.030 (-4.303, 4.363) | 0.701 | -0.869 (-5.333, 3.595) |
| DBP | HannumIEAA | 0.383 | 1.373 (-1.719, 4.466) | 0.273 | 2.419 (-1.921, 6.759) | 0.972 | -0.080 (-4.541, 4.381) |
| DBP | HannumAA | 0.614 | -0.795 (-3.896, 2.306) | 0.721 | -0.786 (-5.121, 3.550) | 0.653 | -1.013 (-5.453, 3.427) |
| DBP | GrimAA | 0.505 | -1.052 (-4.152, 2.047) | 0.039 | -4.530 (-8.832, -0.228) | 0.172 | 3.064 (-1.353, 7.481) |
| DBP | PhenoAA | 0.678 | -0.666 (-3.820, 2.487) | 0.282 | -2.399 (-6.790, 1.992) | 0.547 | 1.390 (-3.173, 5.953) |

## Appendix B. Epigenetic clocks information

## Appendix B.1. First generation clocks

By the first generation clocks we understand epigenetic age predictord which were developed only based on chronological age. Chronological age is in itself a significant risk factor for cardiovascular disease. With age, changes in the cardiovascular system lead to a decline in functioning, and predispose to CVDs such as coronary artery disease (CAD), hypertension, atherosclerosis, stroke, and myocardial infarction (MI). While structural and functional changes which predispose individuals to cardiovascular events (e.g. left ventricular hypertrophy, arrhythmia) have been characterised, the causes of cardiac ageing are not fully understood.

## Appendix B.1.1. Horvath’s clock

This first multi-tissue age estimator was developed on about 8000 samples from 51 different tissues and cell types from both children and adults [3]. A penalised regression model was used to regress transformed chronological age onto the 353 CpGs automatically selected by the elastic net regression model. Specifically, methylation of 193 CpGs positively correlates with chronological age, whilst the other 160 present negative correlation [3]. Together, they form a very accurate molecular measure of chronological age, often used in laboratories for validation of age data of clinical samples [8]. The greatest benefit of Horvath’s clock is the accuracy of age prediction using DNA from a wide range of tissues and organs, with the exception of breast tissue, uterine endometrium, dermal fibroblast, skeletal muscle tissue, heart tissue and sperm cells. Horvath’s age estimate is also reliable in testing all ages, children included [3].

## Appendix B.1.2. Hannum’s clock

Hannum’s clock was developed by regressing chronological age using an elastic net penalised multivariate regression method together with bootstrap approaches which resulted in selection of 71 CpGs as accurate age predictor [2]. This clock was developed on whole-blood DNA from 656 adults leading to possible biases in children and non-blood tissues.

## Appendix B.2. Second generation clocks

The second generation epigenetic clocks are designed to incorporate not only chronological age, but also ageing-related physiological conditions. Specialised clocks (i.e. those which were developed for specific phenotypes) are usually also attributed to the second generation epigenetic clocks.

Intrinsic and extrinsic EAAs With the advance of the second-generation clock, modification to the first generation clocks has proposed to reflect both extrinsic and intrinsic epigenetic age accelerations components (EEAA and IEAA) [65].

IEAA is defined as a residual of the regression of epigenetic age on chronological age and blood immune cell counts, inferred from DNAm data. The list of blood cell types includes naive CD8+ T cells, exhausted CD8+ T cells, plasmablasts, CD4+ T cells, natural killer cells, monocytes, and granulocytes. This EAA measure was created to estimate “pure” epigenetic aging effects that are not influenced by differences in blood cell counts [65].

By design, EEAA is positively correlated with the of counts of exhausted CD8+ T cells, plasmablast cells, and a negative correlated with naive CD8+ T cells, estimated from DNAm data. According to the authors, EEAA was developed to track both age related changes in blood cell composition and intrinsic epigenetic changes[65].

## Appendix B.2.1. Skin and Blood clock

The Skin and Blood Clock’s 391 CpGs were obtained from elastic net regression of chronological age onto CpGs from datasets of human blood, saliva, keratinocytes, buccal cells, endothelial cells, and fibroblasts [29]. In addition to these aforementioned tissues/cell types used in obtaining clock CpGs, Skin and Blood age estimates are strongly correlated with chronological age in other tissues, including colon and heart tissue [29].

Similar to Horvath’s clock, the Skin and Blood clock is accurate across multiple tissues/cell types, and is only weakly affected by blood cell type counts, the clock may capture the physiological processes of cell-intrinsic ageing [8,29]. It may capture cell-intrinsic ageing in cardiac tissues with greater accuracy than Horvath’s clock, since it is capable to detect: (1) age acceleration in individuals with HGPS, who often die from myocardial infarction; and (2) increases in epigenetic age with proliferation of human coronary artery endothelial cells [29].

## Appendix B.2.2. DNAm PhenoAge

Development followed a two stage process. First, a “phenotypic age” metric was created to consider the age-related changes that are associated with senescence, composed from ten clinical characteristics: chronological age, albumin, creatinine, glucose, C-reactive protein, lymphocyte percentage, mean cell volume, red blood cell distribution weight, alkaline phosphatase, and white blood cell count. After that, 513 CpGs were isolated using an elastic net regression of methylation data from whole blood samples onto phenotypic age; this linear combination of CpGs estimates phenotypic age [4].

As DNAm PhenoAge was designed to include CpGs which reflected “phenotypic” ageing rather than chronological ageing [8], it predicts both mortality and age-related morbidity risk with greater accuracy than Horvath and Hannum’s clock [4]. The clinical characteristics used in PhenoAge can be particularly relevant to cardiovascular diseases. For example, albumin is a modulator of vascular functioning and anticoagulation, as well as antioxidation. Creatine has an effect on blood pressure and heart recovery, as well as potentially increasing flow and ATP content while decreasing cell death [66].

## Appendix B.2.3. GrimAge

Development followed a two stage process. First, DNAm-based surrogates for biomarkers of smoking pack-years and twelve plasma proteins were produced, through elastic net regression of DNA methylation data from blood samples. Subsequently, alongside other patient characteristics, the DNAm-based surrogates were considered as covariates in an elastic net Cox regression model for “time-of-death due to all-cause-mortality” [5]. Significant covariates included: chronological age, sex, and DNAm surrogates for smoking pack-years and seven (out of twelve) plasma proteins (ADM, B2M, Cystatin C, Leptin, GDF-15, PAI-1, and TIMP-1). Transformation of the covariates’ linear combination produced the algorithm’s age estimate [5]. According to the authors, GrimAge EAA is a significant predictor of lifespan regardless of smoking history. Furthermore, upon comparison to other clocks (Hannum, Horvath, and DNAm PhenoAge), AgeAccelGrim is a more accurate predictor of age-related disease onset, including coronary heart disease[5].

By design, GrimAge is probably the most relevant epigenetic clock in studying cardiovascular diseases, as the plasma proteins used as physiological variables are known to be associated with CVD. Adrenomedullin (ADM) increases cardiac output and decreases blood pressure [67], while increases of ADM in plasma correspond to hypertension [68]. Beta-2-microglobulin is also an emerging biomarker for cardiovascular diseases [69]. Increased expression of tissue inhibitor of metalloproteinase-1 (TIMP-1) has been linked to cardiac fibrosis and dysfunction of the heart [70]. Given that a significant number of the physiological variables used in the development of GrimAge have been shown to associate strongly specifically with cardiovascular pathology, it is important to consider this clock in the study of cardiovascular disease.

## Appendix B.3. Summary of the epigenetic clocks

Epigenetic clocks use mathematical modelling to evaluate the methylation of the specific CpG sites in DNA in order to estimate the epigenetic age. While the epigenetic age highly correlates with the chronological age, a difference between chronological and epigenetics age, termed the epigenetic age acceleration (EAA) can be observed and has been studied extensively. EAA has been associated with many later-life pathologies ranging from cardiovascular and neurological diseases to cancer. To reflect the changes in celltype composition of blood naturally occurring with ageing, the extrinsic epigenetic age acceleration (EEAA) has been termed. These changes are more accurately measured by clocks from blood DNA, such as Hannum’s or PhenoAge [8]. On the other hand, the cell-intrinsic ageing, which is believed to be more consistent over various tissues and organs is more accurately measured by the Horvath’s multi-tissue clock, and has been termed the intrinsic epigenetic age acceleration (IEAA) [8].

Epigenetic clocks use different CpGs, and therefore naturally form different associations with different disease processes, and the ways in which the clocks are constructed is highly relevant to the processes they capture. Specifically, associations between certain CpGs and the biological processes underlying development, maintenance of cellular identity and cell differentiation have been captured [8]. Another possible association has been made with regards to the circadian rhythm, suggesting that epigenetic ageing might be linked to genetic oscillation, such as cell cycle oscillator. One review compared six different types of ageing biomarkers, and found that epigenetic clocks were the best predictors of chronological age.

Both first generation clocks used in our study (Horvath and Hannum) were constructed to accurately predict chronological age. PhenoAge and GrimAge both include a training step which uses physiological variables, which may explain why both of these clocks appear to be more sensitive to age-related pathologies than first generation clocks constructed for the sole purpose of predicting chronological age.

For example, smoking, a known risk factor for disease, is not reflected in the Horvath and Hannum clocks, despite being a strong DNAm mortality predictor [71]. PhenoAge captures smoking-associated methylation changes, and GrimAge uses a DNAm surrogate of pack-years, both of which lead to a better prediction of smoking-related pathology [5].

It is not entirely clear what is measured by epigenetic clocks and to what extent biological ageing is related to disease. In the review [72] the authors suggest that epigenetic clocks track both CA and physiological/pathological mechanisms to differing proportions.

## Notes

### Competing Interest Statement

The authors have declared no competing interest.

### Summary of Updates

We added extended supplementary by adding a table with all the testing results, along with an overview summary on the epigenetic clock models. There are also some minor text edits throughout the manuscript.

